# Directed evolution of compact RNA-guided nucleases for enhanced activity in mammalian cells

**DOI:** 10.1101/2025.10.27.684765

**Authors:** Fedor Gorbenko, Irene Sala, Young-Yoon Lee, Lilly van de Venn, Charles D Yeh, András Tálas, Tautvydas Karvelis, Gytis Druteika, Luca V. Bechter, Iryna Vykhlyantseva, Markus S. Schröder, Ana Gvozdenovic, Gerald Schwank, Virginijus Siksnys, Jacob E. Corn

## Abstract

RNA-guided nucleases enable DNA editing and offer promise for treating genetic diseases, particularly when used for precise sequence replacement. However, many of the most effective enzymes, such as *S. pyogenes* Cas9, are too large for delivery using vectors like adeno-associated virus (AAV). This has prompted interest in smaller alternatives from the Cas12f and TnpB families. Yet, these nucleases often show low activity in mammalian cells, limiting their utility. To address this, we used directed evolution in human cells to select variants with greatly improved activity. The resulting variants, Cas12f1Super and TnpBSuper, exhibited up to 11-fold increase in editing efficiency without increased off-target effects. When tested as a base editor, Cas12f1Super showed up to 10-fold improvement relative to the previously engineered CasMINI, suggesting utility beyond nuclease-related activities. These compact and efficient genome editors expand the current toolkit and hold promise for both research and therapeutic use in mammalian systems.

## Introduction

CRISPR-Cas technology has revolutionized genome editing by enabling precise, RNA-guided manipulation of genomic DNA in a wide range of contexts. This advancement has unlocked powerful applications in biomedical research and holds great promise for treating genetic diseases through targeted genome correction. However, most existing highly active CRISPR systems, such as Cas9 and Cas12a, are large proteins, limiting their compatibility with clinically relevant size-restricted delivery vectors like adeno-associated viruses (AAVs)^1^.

This constraint has driven growing interest in the development of compact genome editing systems that are more compatible with AAVs and other delivery platforms. Several miniature nucleases, including Cas12f and TnpB-based systems, have recently been explored as alternatives^2^. However, these proteins exhibit substantially lower editing efficiencies in mammalian systems than larger Cas enzymes, limiting their applicability and stimulating efforts to improve their activity^3–9^.

Efforts to enhance enzyme activity typically rely on rational design or directed evolution. While rational design leverages structural and mechanistic insights to introduce beneficial mutations, it is inherently limited by our incomplete understanding of protein function and may miss unexpected improvements. Directed evolution allows broader exploration through functional selection from diverse variant pools. It is often performed in non-human systems, such as phage or bacteria^9,10^, which offers several technical advantages such as the ability to interrogate large libraries necessary to cover large Cas proteins. However, these hosts fail to recapitulate many aspects of human cell biology, ranging from intracellular folding propensity to chromatin structure. Their DNA repair responses can also differ greatly from human cells, which for genome editing purposes necessitates evolution towards proxy targets such as targeting binding or catalytic activity. This may yield variants with high activity in the evolved context but modest improvement when applied to genome editing in human cells.

The effectiveness of directed evolution in mammalian systems is constrained by the scale of cell culture. For example, screening 2.5 × 10^7^ variants in mammalian cells would allow testing of only about 7% of all possible single and double amino acid mutants for SpCas9 (1368 amino acids). This consideration underscores another advantage of compact editors. With only 528 and 408 amino acids, respectively, Un1Cas12f1 and ISDra2 TnpB require far fewer variants to cover a meaningful fraction of their sequence space^11,12,13^. Screening 2.5 × 10^7^ variants covers approximately 50% of all single and double mutants for Un1Cas12f1 and 83% for ISDra2 TnpB. This reduced search space allows for effective directed evolution of desired genome editing activities in mammalian cells. While all possible single point mutations of AsCas12f1 have recently been tested by deep mutational scanning (DMS), potentially more active variants with multiple mutations have never been explicitly selected in human cells via directed evolution^8^.

For many genome editing applications, including therapeutic gene correction, nuclease activity alone is not sufficient. Editing is the result of host cell repair processes, including high-fidelity homology-directed repair (HDR) that enables precise sequence changes when a donor template is present. In mammalian cells, HDR is usually outstripped by the faster, error-prone non-homologous end joining (NHEJ) pathway, leading to several workaround technologies such as base and prime editing^14,15^. But base and prime editing add further size to the editing architecture, again emphasizing the importance of compact editors. Improving editing efficiency, particularly precise sequence manipulation such as HDR, remains a major obstacle to expanding the utility of small genome editors for clinical use. In this regard, mammalian-based selection offers a key advantage: it allows identification of variants that specifically enhance editing performance in therapeutic-relevant settings rather than catalytic activity.

Here we performed mammalian cell directed evolution on two small genome editing proteins: Un1Cas12f1, a miniature Cas12f1-family nuclease derived from archaea, and ISDra2 TnpB, a TnpB-family nuclease derived from bacteria. We applied directed evolution in a fluorescent reporter system to isolate variants with increased HDR efficiency. Through iterative rounds of mutagenesis and screening, we assembled combinatorial variants that show robust HDR improvements achieving up to 11-fold increase in editing efficiency. The resulting variants also exhibit increased NHEJ activity. We also demonstrated the portability of the best Un1Cas12f1 variant by turning it into an adenine base editor with up to a 10-fold higher editing efficiency relative to the engineered Un1Cas12f1 baseline. The engineered “Cas12f1Super” and “TnpBSuper” enzymes reported here represent promising tools for efficient genome editing in mammalian systems for both fundamental research and therapeutic applications.

## Results

### Directed evolution of Un1Cas12f1 and ISDra2 TnpB for enhanced activity in mammalian cells

To enable directed evolution of compact nucleases, we first developed a fluorescent HDR reporter system for selecting, screening, and quantifying HDR-active variants in mammalian cells (**Fig. 1a**). This reporter contained a disrupted start codon in the[*EGFP*[gene, making *EGFP* expression dependent on successful HDR-mediated repair. A customizable protospacer and PAM/TAM directly upstream of[*EGFP*[allow precise targeting by various single guide RNAs (sgRNAs) or ωRNAs and assessment of HDR activity across different target sequences. These reporters (one for Un1Cas12f1 and one for ISDra2 TnpB) were polyclonally integrated into HEK293T cells at moderate multiplicity of infection, selecting for blasticidin resistance present in the cassette. For ease of recognition, we refer to these protospacer/PAM/TAM combinations as “Target CM1” for Un1Cas12f1 and “Target TB1” for ISDra2 TnpB (see Methods).

**Figure 1.**
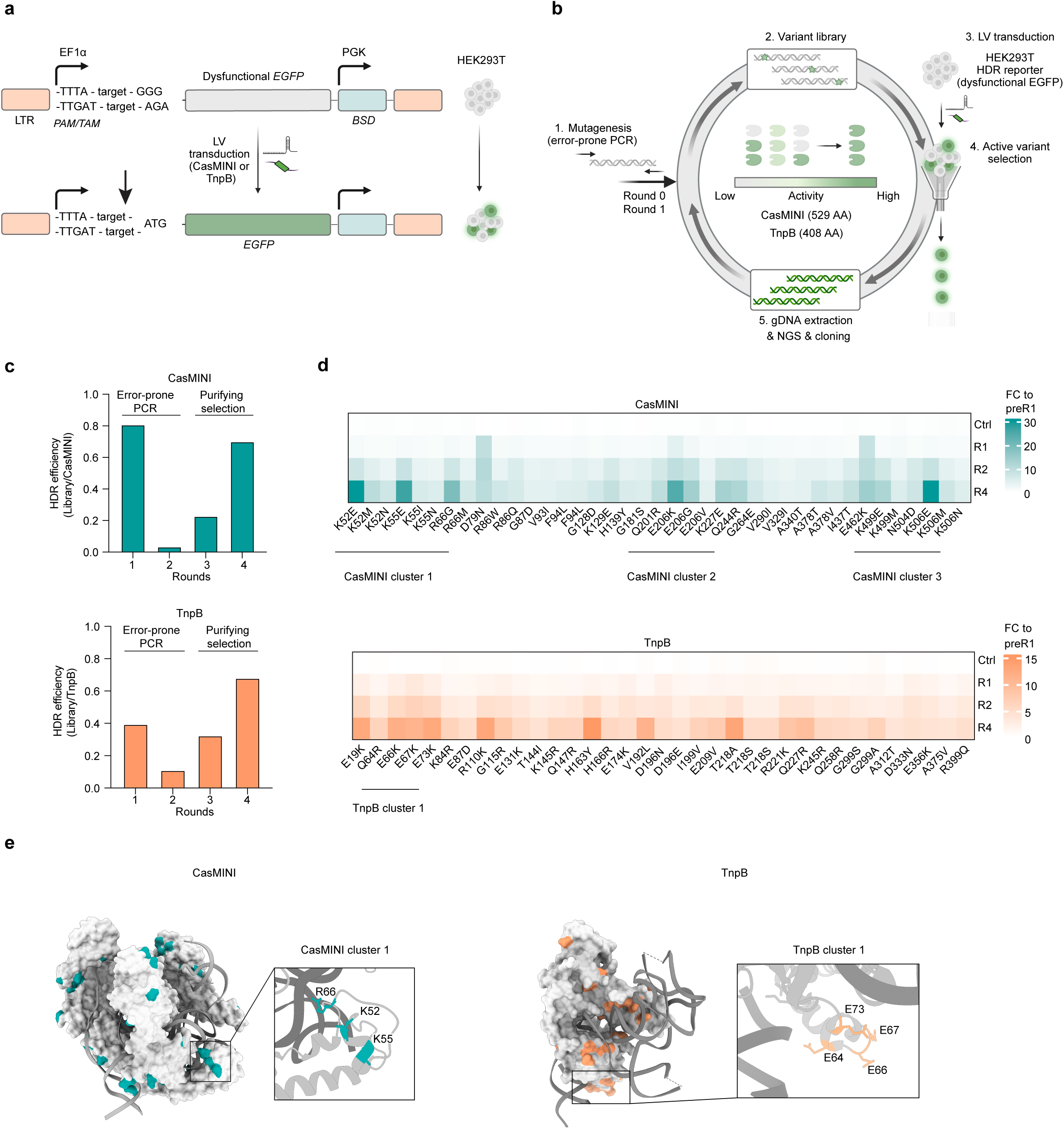
Directed evolution of CasMINI and TnpB for enhanced HDR activity in mammalian cells. (**a**) Schematic of the homology-directed repair (HDR) fluorescent reporter. A protospacer with its protospacer-associated motif (Un1Cas12f1 PAM: TTTA) or transposon-associated motif (ISDra2 TnpB TAM: TTGAT) was cloned upstream of an *EGFP* gene lacking a start codon. Lentiviral delivery into HEK293T cells generated a non-fluorescent polyclonal cell line. *EGFP* expression can be restored upon successful HDR following delivery of a nuclease (CasMINI or TnpB), a guide RNA (sgRNA for CasMINI and ωRNA for TnpB), and a donor single-stranded oligodeoxynucleotide (ssODN) containing the start codon. An arrow indicates approximate cut position. *BSD*-gene for blasticidin resistance, LTR-long terminal repeat; LV-lentiviral. (**b**) Schematic of the directed evolution selection workflow. CasMINI or TnpB variant libraries were screened in reporter cells by iterative cycles of HDR, fluorescence-activated cell sorting (FACS) enrichment, and library regeneration to select for improved activity. (**c**) HDR efficiency measured as bulk EGFP signal across four rounds of selection of indicated proteins, normalized to the respective wild-type nuclease. Bars represent the value of a single experimental replicate. (**d**) Relative frequency of enriched mutations of the original sequences (Ctrl) and after rounds 1, 2, and 4 (“R1”, “R2”, “R4”, respectively) of selection for indicated nucleases. Mutations with >0.3% frequency and >3.75-fold enrichment over the pre-selection libraries are shown. The clusters with enriched mutations are marked. FC, fold-change. (**e**) Enriched mutations mapped onto the structures of nuclease Un1Ca12f1 (PDB: 7l49), highlighted in teal and nuclease TnpB (PDB: 8H1J) highlighted in orange. Close-up views for mutation cluster 1 are presented for each nuclease. Protein structures were visualized with ChimeraX^45^.

To evaluate the baseline HDR activity of Un1Cas12f1 and ISDra2 TnpB proteins in this format, we delivered each nuclease by lentiviral transduction with puromycin selection and subsequently electroporated both a plasmid expressing guide RNA (sgRNA for Un1Cas12f1 and ωRNA for TnpB) and a single-stranded oligodeoxynucleotide (ssODN) donor template for *EGFP* sequence repair. The percentage of EGFP-positive cells served as a readout of HDR efficiency. ISDra2 TnpB was tested in its wild-type (WT) form, hereafter referred to as TnpB. For Un1Cas12f1, we tested a recently engineered version that combines CasMINI V3.1^3^ with the ge4.1 guide scaffold^6^, both of which were previously shown to improve activity. For simplicity, we hereafter refer to this combination as CasMINI. Both CasMINI and TnpB exhibited detectable HDR at their respective targets (Target CM1 or Target TB1) as measured by EGFP-positive cell percentage (CasMINI: 16.1±3.6% and TnpB: 6.3±0.5%) only once both sg/ωRNA and ssODN were delivered (**Fig. S1a**). HDR alleles were verified for both systems by amplicon next-generation sequencing (ampliconNGS) (**Fig. S1b**). This established a foundation for using this reporter system for the selection of activity-enhancing mutations.

To perform directed evolution, we first performed error-prone polymerase chain reaction (epPCR) to generate libraries of 1.5 × 10^7^ variants for CasMINI and 4.5 × 10^7^ variants for TnpB, which is an average of ∼2 amino acid–altering mutations per enzyme. We deliberately chose a low mutagenesis rate to avoid the accumulation of deleterious changes that could mask beneficial effects. As a result, about half of each library remained unmutated WT sequence (**Fig. S2a**). Each library was delivered to cells harboring its corresponding Target CM1 or Target TB1 within the *EGFP* reporter system. Cells were electroporated with a plasmid expressing sg/ωRNA targeting the reporter along with ssODN donor to restore *EGFP* expression. EGFP-positive cells were isolated by fluorescence-activated cell sorting (FACS), and their genomic DNA was used for targeted PCR amplification of active variants for cloning into a new plasmid library for the next round of selection (**Fig. 1b**).

We performed two initial rounds of selection with intervening error-prone PCR to increase library diversity (**Fig. 1b**). Bulk library activity dropped during these rounds, consistent with early rounds of selection that had not yet enriched highly active variants (**Fig. 1c**). We then carried out two additional rounds of purifying selection without further mutagenesis, during which EGFP levels gradually increased, indicating an enrichment for more active variants.

After rounds 1, 2, and 4 of selection we amplified and tagmented the nuclease construct and used NGS to determine the unphased mutation spectrum of the bulk pool. For both CasMINI and TnpB we found multiple amino acid changes that were steadily enriched compared to the pre-selection libraries (**Fig. 1d**). In case of CasMINI we identified three clusters of enriched mutations (clusters 1-3). Among them, CasMINI cluster 1 exhibited enrichment of mutations at spatially proximal residues 52, 55, 66 (**Fig. 1d,e, S2b**), which are located near or within hydrogen-bonding distance to the sgRNA. Additionally, we observed two clusters (residues 201, 206, 227, 244 and 499, 504, 506) containing at least one highly enriched mutation (**Fig. 1d, S2b**). Among the 39 most enriched mutations in CasMINI, 28 involved a change in charge. Surprisingly for a DNA-binding protein, 24 of these selected mutations lost positive charge or gained negative charge, including the most abundantly selected K52E, K55E, and K506E mutations. E206K was also a highly enriched mutation, which is similar to the previously reported E206R^16^.

The selected TnpB pool exhibited a broader distribution of enriched mutations with only one obvious cluster at positions 64, 66, 67, 73 (TnpB cluster 1), which the structure of TnpB revealed to be in proximity to DNA (**Fig. 1d,e, S2c**). For TnpB, 17/34 most enriched mutations involved a change in charge. In contrast to what we observed for CasMINI, only one of these showed a loss of positive charge. The rest exhibited a loss of negative charge or a gain of positive charge. This included mutations from TnpB cluster 1: E64K, E66K, E67K, and E73K. Overall, enrichment of multiple mutations for both nucleases after selection for HDR prompted us to investigate these mutations as potential drivers of HDR activity.

### Generation of CasMINI and TnpB with enhanced activity

The unphased short-read sequencing we used to measure mutation abundance in the selected pools cannot resolve co-occurring beneficial mutations. To develop high-performing CasMINI and TnpB variants, we isolated and tested promising multi-mutation variants from *in cellulo* evolution. Mutations from the best variants were combined and tested at multiple reporter and endogenous targets (**Fig. 2a**). This strategy provided a structured path from a diverse variant pool to functionally robust and generalizable genome editing configurations.

**Figure 2.**
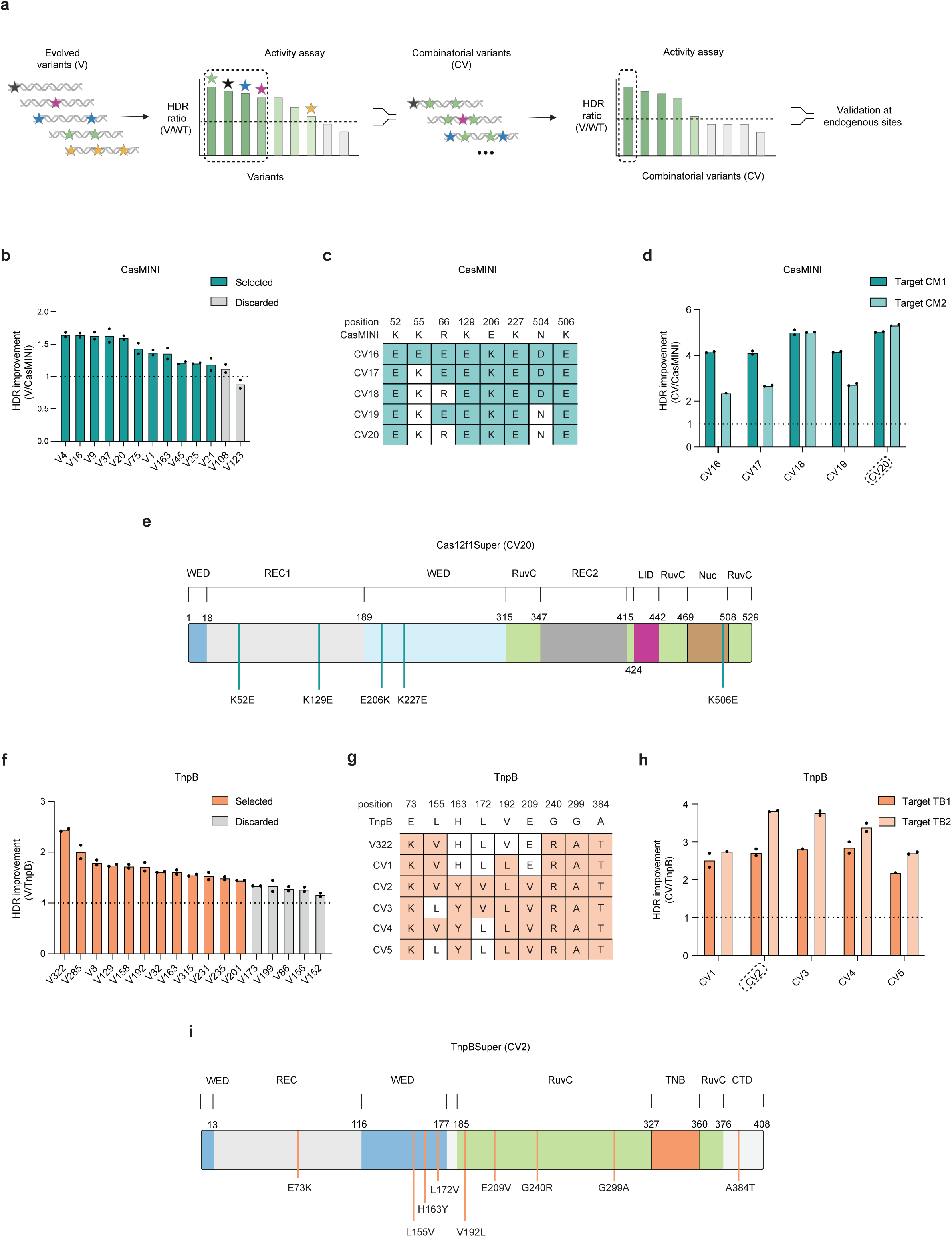
Generation of CasMINI and TnpB variants with enhanced activity. (**a**) Schematic of the strategy for generating combinatorial variants (CVs). Screen-identified variants were ranked by homology-directed repair (HDR) activity and used to guide combinatorial assembly. An additional HDR reporter target was used to exclude target-specific effects. Final combinatorial variants were validated at endogenous genomic sites. (**b**) HDR activity of post-round 4 variants measured as percentage of EGFP-positive cells from BFP/mCherry double positive cells (BFP marks the guide-expressing construct; mCherry marks the nuclease-expressing construct), normalized to CasMINI (*n* = 2 biological replicates). Variants exceeding 1.2-fold, highlighted in teal, were used to identify active mutations. (**c**) A table showing mutations present in CasMINI combinatorial variants 16-20 (CVs 16-20). (**d**) HDR activity of CVs at two targets measured as percentage of EGFP-positive cells from BFP/mCherry double positive cells (BFP marks the guide-expressing construct; mCherry marks the nuclease-expressing construct), normalized to CasMINI (*n* = 2 biological replicates). The CV selected for further testing (CV20) at endogenous sites is highlighted. (**e**) Mutations present in Cas12f1Super (CV20) are mapped onto the domain organization of Un1Cas12f1 (PDB: 7l49). (**f**) HDR activity of post-round 4 variants measured as percentage of EGFP-positive cells from BFP/mCherry double positive cells (BFP marks the guide-expressing construct; mCherry marks the nuclease-expressing construct), normalized to TnpB (*n* = 2 biological replicates). Variants exceeding 1.4-fold, highlighted in orange, were used to identify active mutations. (**g**) A table showing mutations present in V322 and combinatorial variants 1-5 (CVs 1-5) of TnpB. (**h**) HDR activity of CVs at two targets measured as percentage of EGFP-positive cells from BFP/mCherry double positive cells (BFP marks the guide-expressing construct; mCherry marks the nuclease-expressing construct), normalized to TnpB (*n* = 2 biological replicates). The CV selected (CV2) for further testing at endogenous sites is highlighted. (**i**) Mutations present in TnpBSuper (CV2) are mapped onto the domain organization of TnpB (PDB: 8H1J). Each dot represents an individual biological replicate, and bars represent the mean.

For CasMINI, we tested a total of 246 post-round four variants in HEK293T cells (88 tested with stable expression using lentiviral delivery and 158 transiently expressed from plasmids, **Fig. S3a**). We first tested these variants against the same target site used for selection (Target CM1), resulting in eleven multi-mutation variants with a 1.2 to 1.6-fold increase in HDR efficiency relative to CasMINI (**Fig. 2b, Supplementary Table 1**). To isolate activity drivers from passengers, we cloned and tested 22 mutations from the promising variants as single mutants (SMs) in the CasMINI background. We also tested 18 additional candidate SMs identified as enriched by NGS analysis of the selected pool (**Fig. 1d**), resulting in a total of 40 tested mutations (**Fig. S3b**). We retested the top-performing 19 mutations against an orthogonal target site (hereafter “Target CM2”, see Methods) and identified eight mutations that consistently enhanced HDR at both targets (**Fig. S3c**). We combined eight selected mutations with high activity at both target sites into five distinct combinatorial variants (CVs) (**Fig. 2c, Supplementary Table 1**). Rather than performing an exhaustive combinatorial screen, we employed a structure-guided reduction strategy in which we cloned a “master” variant with all selected mutations that individually enhanced activity and then derived sub-variants by selectively reverting subsets of mutations to the parental amino acid. The design was guided by structural proximity of mutations, with the aim of reducing the likelihood of negative epistatic interactions between closely positioned residues. In individual testing using plasmid expression, all CVs significantly outperformed both CasMINI and the individual mutations at two independent target sites. Comparison of CV16, CV17 and CV18 showed that having K52E alone (CV18) is more beneficial than having either K52E, K55E, R66E altogether (CV16) or K52E paired with R66E (CV17). CV18 (six mutations) and CV20 (five mutations) showed the highest levels of HDR, with CV20 achieving 5.3-fold improvement relative to CasMINI at Target CM2 (**Fig. 2d**).

We selected CV20 for further studies (**Fig. 2d**) based on its high expression in HEK293T cells (**Fig. S3d**) and target-independent activity. CV20 contains five additional mutations on top of CasMINI (K52E, K129E, E206K, K227E, K506E), with at least one mutation from each of the three clusters that we previously identified that map to the REC, WED and NUC domains of CasMINI (**Fig. 2e**). We attempted to further improve CasMINI_CV20 using the recently described EVOLVEpro machine learning framework^17^, training on activity data collected during single mutant testing (**Fig. S3c**). Eight different EVOLVEpro-suggested point mutations introduced in the CV20 revealed no further improvements in activity (**Fig. S3e**), and so we continued with CV20.

To assess the portability and contribution of the evolved mutations, we incorporated the CV20 mutations into the WT Un1Cas12f1 background (Un1Cas12f1_CV20) instead of the CasMINI background. We then compared the HDR activities of WT Un1Cas12f1, Un1Cas12f1_CV20, CasMINI and CasMINI_CV20 at both Target CM1 and Target CM2. We observed that the CV20 mutations greatly increased the activity of both WT Un1Cas12f1 and CasMINI (**Fig. S3f**). The HDR activity of CasMINI_CV20 was up to 11.4±1.2-fold greater than that of WT Un1Cas12f1, so we selected it for later testing at endogenous sites. We termed this final system with five coding mutations relative to CasMINI as Cas12f1Super **(Fig. 2e)**.

For TnpB, we tested a total of 311 post-round four variants in HEK293T cells (**Fig. S4a**, all tested using plasmid expression). This yielded 12 variants with at least 1.4-fold improvement (**Fig. 2f, Supplementary Table 1**) at the Target TB1 site used for selection. One variant, designated V322 (five amino acid altering mutations) consistently exhibited the highest HDR activity, achieving 2.4-fold improvement over TnpB. Notably, V322 contained E73K, which was the most enriched mutation within the TnpB cluster 1 after four rounds of selection (**Fig. 1d**). To expand our search for stronger contributors, we cloned and tested 17 mutations from the 12 TnpB variants, as well as 19 additional mutations identified by NGS of the selected pool resulting in total of 36 SMs. In total, we identified 16 constructs that improved HDR by at least a 1.9-fold over TnpB (**Fig. S4b**). We retested the top-performing variant and single mutations at an orthogonal target (hereafter “Target TB2”, see Methods) (**Fig S4c**). Several SMs performed well at both target sites, and we selected a subset of these broadly effective substitutions to incorporate on top of V322, yielding five CVs (**Fig. 2g, Supplementary Table 1**). As with CasMINI, we sought to limit combining proximal mutations within the same variant, based on the assumption that their effects might be redundant or negatively epistatic. All CVs exhibited at least two-fold greater HDR than WT TnpB at both Target TB1 and Target TB2. We chose to move forward with the high-performing CV2 (**Fig. 2h**), which carried nine mutations (**Fig. 2i**). We tested eight additional EVOLVEpro-suggested mutations in the TnpB_CV2 background (**Fig. S4d**)^17^, but none strongly increased activity.

Two final refinements of TnpB_CV2 and ωRNA further improved activity and were incorporated into the final design. While CasMINI (and hence Cas12f1Super) was already codon-optimized for mammalian cells, we performed selection with TnpB cloned with its original bacterial codon usage. Mammalian codon optimization slightly improved the activity of both TnpB and TnpB_CV2 across two targets (**Fig. S5a,b**). However, we observed no great increase in the expression of TnpB_CV2 compared to TnpB in HEK293T cells **(Fig. S5c)**. A recently reported shorter version of the ωRNA (Trim2)^18^ also marginally improved activity for TnpB_CV2 at both target sites (**Fig. S5d,e**). We next explored an additional refinement by combining the TnpBSuper mutations with the previously developed TnpBmax construct design^19^ to generate TnpBmaxSuper. However, the addition of TnpBmax architecture did not improve activity at either of the two HDR reporters nor at two endogenous sites (**Fig. S5f,g**), which is an unexpected result in light of prior observations with TnpBmax as a nuclease^19^. We therefore did not incorporate this optimization in subsequent experiments. We termed the final codon-optimized system with nine coding mutations relative to TnpB and the Trim2 ωRNA as TnpBSuper **(Fig. 2i)**.

### Cas12f1Super and TnpBSuper efficiently edit endogenous genomic loci

To evaluate whether the observed improvement in HDR for Cas12f1Super and TnpBSuper translated to endogenous gene targets, we edited 13 distinct genomic sites in HEK293T cells using plasmid-based delivery of nuclease and sg/ωRNA (**Fig. 3a**). Editing outcomes were measured using ampliconNGS after enriching for both guide RNA and editor-positive cells by FACS.

**Figure 3.**
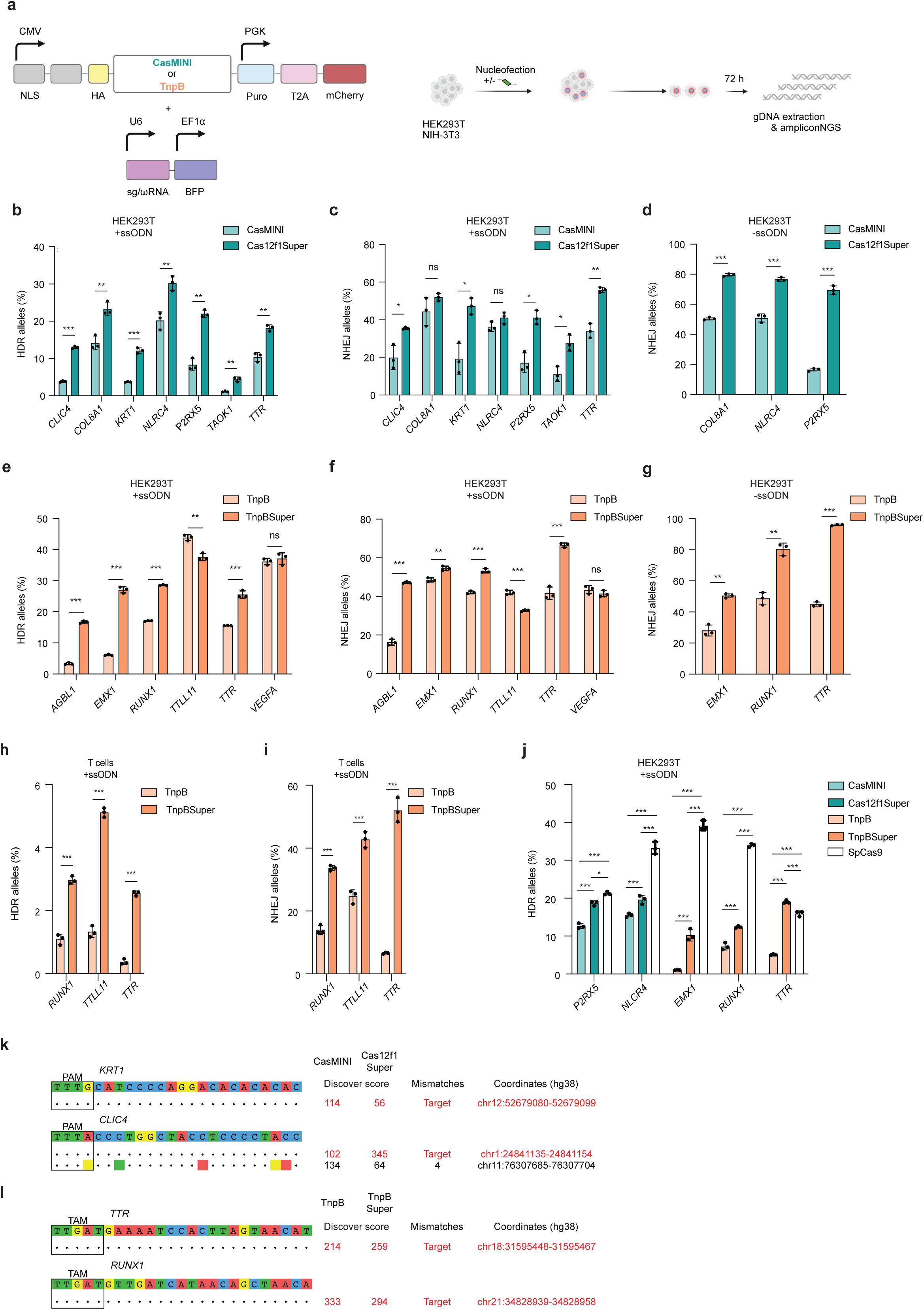
Improved variants Cas12f1Super and TnpBSuper edit endogenous genomic loci. (**a**) Schematic of the workflow of endogenous site editing in HEK293T and NIH-3T3 cells. Nuclease-expressing plasmid (containing mCherry marker) was co-delivered with guide-expressing plasmid (containing BFP marker). ssODN was used only for HDR-related experiments in HEK293T cells. After enrichment for cells that had taken up both plasmids, ampliconNGS was performed. (**b,c**) Absolute HDR (**b**) or NHEJ efficiency of Cas12f1Super (**c**) in the presence of ssODN measured with ampliconNGS at indicated endogenous loci in HEK293T cells compared to CasMINI (*n* = 3 biological replicates). (**d**) Absolute NHEJ efficiency in the absence of ssODN of Cas12f1Super measured with ampliconNGS at indicated endogenous loci in HEK293T cells compared to CasMINI (*n* = 3 biological replicates). (**e,f**) Absolute HDR (**e**) or NHEJ efficiency of TnpBSuper (**f**) in the presence of ssODN at indicated endogenous loci in HEK293T cells compared to TnpB measured with ampliconNGS (*n* = 3 biological replicates). (**g**) Absolute NHEJ efficiency in the absence of ssODN of TnpBSuper measured with ampliconNGS at indicated endogenous loci in HEK293T cells compared to TnpB (*n* = 3 biological replicates). (**h,i**) Absolute HDR (**h**) or NHEJ efficiency (**i**) of TnpBSuper at indicated endogenous loci in primary T cells compared to TnpB measured with ampliconNGS (*n* = 3 biological replicates). TnpBSuper or TnpB were electroporated as mRNA, along with synthetic ωRNA and ssODN. (**j**) Comparison of CasMINI/Cas12f1Super and TnpB/TnpBSuper with SpCas9 measured with ampliconNGS at indicated endogenous loci in HEK293T cells (*n* = 3 biological replicates). (**k,l**) Off-target sites identified by DISCOVER-Seq. Sequences are presented along with Discover-seq scores for on-target and off-target editing with CasMINI/Cas12f1Super (**k**) or TnpB/TnpBSuper **(l)**. Each dot represents an individual biological replicate, and bars represent the mean ± standard deviation. All *P* values were calculated using an unpaired, two-sided *t*-test. ns, not significant (P ≥ 0.05), \**P*L<L0.05, \*\**P*L<L0.01, \*\*\**P*L<L0.001.

For Cas12f1Super we observed consistent and significant increases in HDR efficiency across all seven sites (**Fig. 3b**), with improvements up to 4.0±0.8-fold relative to CasMINI at *TAOK1* **(Fig. S6a**) At one target site, *NLRC4*, absolute HDR efficiency reached up to 30±2% (**Fig. 3b)**. NHEJ rates were increased at five of seven targeted sites, with 2.5±1.0%-fold improvement at *TAOK1* (**Fig S6b**) and reaching up to 56±1.4% at *TTR* (**Fig. 3c**). Next, we evaluated the activity of Cas12f1Super in the absence of an ssODN donor to determine whether enhanced activity would also be observed under these conditions. Across all three endogenous sites tested, Cas12f1Super exhibited increased editing efficiency (**Fig. 3d**), with improvements of up to 4.2±0.3-fold at *P2RX5* (**Fig. S6c**). At the *COL8A1* site, absolute NHEJ editing efficiency reached 80±0.8% (**Fig. 3d**). Together, these results show that Cas12f1Super is generally more active than CasMINI in mammalian cells for both HDR and NHEJ outcomes.

Intrigued by the enhancement in activity of the evolved proteins, we compared the *in vitro* activity of CasMINI and Cas12f1Super. We expressed and purified both nucleases and tested cleavage at six different temperatures (ranging from 37°C to 55°C) at two targets (*KRT1* and Target CM1) **(Fig. S7a-f)**. We observed no significant differences in cleavage activity between CasMINI and Cas12f1Super at any of the temperatures. The cleavage pattern also remained the same in the evolved variant (**Fig. S7g,h).** The improvement in HDR and NHEJ for CasMINI may therefore be unrelated to catalytic activity in the tube. The high activity of these variants could instead reflect interactions with the mammalian cell environment, though this remains to be tested and may be difficult to determine.

TnpBSuper demonstrated significant increases in HDR at four of six endogenous sites (**Fig. 3e**), with HDR improvements up to 5.0±0.5-fold over TnpB at *AGBL1* (Fig. S8a). NHEJ efficiency was increased relative to TnpB at the same four sites (**Fig. 3f, S8b**). We next assessed the activity of TnpBSuper in the absence of an ssODN donor, analogous to the analysis performed for CasMINI and Cas12f1Super. Across all three endogenous sites tested, TnpBSuper exhibited increased editing efficiency (**Fig. 3g**), with improvements of up to 2.1±0.1-fold at *TTR* (**Fig. S8c**) and absolute editing efficiency reaching 96±0.5% at the same site (**Fig. 3g**). Overall, these results show that TnpBSuper outperforms TnpB in mammalian cells across both HDR and NHEJ.

In order to further explore the potential of enhanced cutting efficiency of Cas12f1Super and TnpBSuper, we tested all possible guides targeting exon 1 of the *TRAC* locus in HEK293T cells, including nine guides for CasMINI and one guide for TnpB. For the majority of guides, we observed efficient editing, with higher activity for the evolved variants compared to their respective wild-type enzymes (**Fig. S8d**). Together, these results further support the enhanced activity of the evolved systems across diverse guide sequences.

We next tested activity in the clinically relevant setting of primary human T cells. We produced mRNAs for TnpB and TnpBSuper and co-electroporated them with synthetic ωRNA and an appropriate ssODNs targeting three endogenous loci in activated primary T cells. TnpBSuper exhibited consistent increases in HDR relative to TnpB at all three sites, with improvements reaching up to 6.9±1.6-fold at *TTR* (**Fig. S8e**), and absolute HDR efficiencies up to 5.1±0.1% at *TTLL11* (**Fig. 3h**). Enhancements in NHEJ were also significant, with up to 7.9±0.6-fold improvement and levels reaching 52±4% at the *TTR* site (**Fig. S8f, 3i**).

To benchmark our evolved systems, we compared their performance to that of the widely used but much larger *S. pyogenes* Cas9 (*Sp*Cas9) nuclease. We selected two loci for CasMINI system (*P2RX5* and *NLRC4*) and three loci for TnpB system (*EMX1*, *RUNX1*, and *TTR*) and identified up to three SpCas9 guides per locus based on close proximity to CasMINI/TnpB target sites and high Rule Set1 activity scores^20^. These guides were first evaluated for NHEJ activity in mammalian cells (**Fig. S8g**), and the best-performing guide at each locus was selected for direct comparison. We then compared SpCas9 side by side with CasMINI/Cas12f1Super or TnpB/TnpBSuper for HDR efficiency (**Fig. 3j**). SpCas9 outperformed Cas12f1Super and TnpBSuper at most loci, consistent with over ten years of development as a workhorse genome editing system. However, at the *TTR* site, TnpBSuper achieved slightly higher HDR (19±0.4%) compared to SpCas9 (16±0.7%). At the *P2RX5* locus, Cas12f1Super approached SpCas9 performance, with HDR efficiencies of 19±0.6% and 21±0.5%, respectively. Cas12f1Super and TnpBSuper outperformed their parental constructs in all cases.

The enhanced genome editing activity of Cas12f1Super and TnpBSuper could potentially come at the cost of increased off-target effects. To evaluate this possibility, we employed DISCOVER-Seq, which enables nuclease-agnostic off-target identification in mammalian cells by monitoring the recruitment of DNA repair factors^21^. For each nuclease pair (CasMINI/Cas12f1Super and TnpB/TnpBSuper), we tested two target sites: *KRT1* and *CLIC4* for CasMINI/Cas12f1Super (**Fig. 3b**), and *TTR* and *RUNX1* for TnpB/TnpBSuper (**Fig. 3e**). Consistent with the previously reported high fidelity of both CasMINI and TnpB, we observed only a single off-target site for CasMINI at the *CLIC4* locus. NHEJ efficiency at this off-target was comparable between CasMINI and Cas12f1Super (**Fig. 3k,l, Fig. S9a,b**). Importantly, despite their enhanced on-target activity, neither Cas12f1Super nor TnpBSuper exhibited additional off-target effects. These results demonstrate that the evolved, highly active variants retain the high specificity of their parental nucleases.

### Efficient in vivo editing using AAV delivery of Cas12f1Super and TnpBSuper

Encouraged by the performance of Cas12f1Super and TnpBSuper at endogenous loci in HEK293T and T cells, we further tested these variants *in vivo* in the context of a therapeutically relevant target gene. We selected *PCSK9*, which is an attractive target for *in vivo* AAV delivery that would be enabled by the hyper-active miniature genome editors developed here^22,23^. *PCSK9* encodes a liver-expressed protein that promotes degradation of the low-density lipoprotein receptor (LDLR). Disrupting PCSK9’s function leads to reduced levels of circulating low-density lipoprotein cholesterol (LDL-C), a strategy that can be exploited for treating familial hypercholesterolemia^24^. Since PCSK9 loss-of-function is sufficient to reduce LDL-C, we leveraged the increased NHEJ activity we observed for Cas12f1Super and TnpBSuper. We first screened for sgRNAs and ωRNAs targeting exons 2-9 of *Pcsk9* in murine NIH-3T3 cells and identified five top-performing guides: three for Cas12f1Super and two for TnpBSuper (**Fig. 4a,b**). These guides were then used to compare the editing efficiency of *Pcsk9* by CasMINI, Cas12f1Super, TnpB, and TnpBSuper in the murine hepatoma cell line Hepa 1-6. We found that Cas12f1Super outperformed CasMINI at all three sites and by up to 2.0±0.13-fold at site 3 (**Fig. S10a**) and reaching 51±1.6% absolute efficiency at site 7 (**Fig. 4c**). TnpBSuper outperformed TnpB at site 4 by 3.2±0.5-fold (**Fig. S10b**), reaching 43±6% absolute efficiency (**Fig. 4d**).

**Figure 4.**
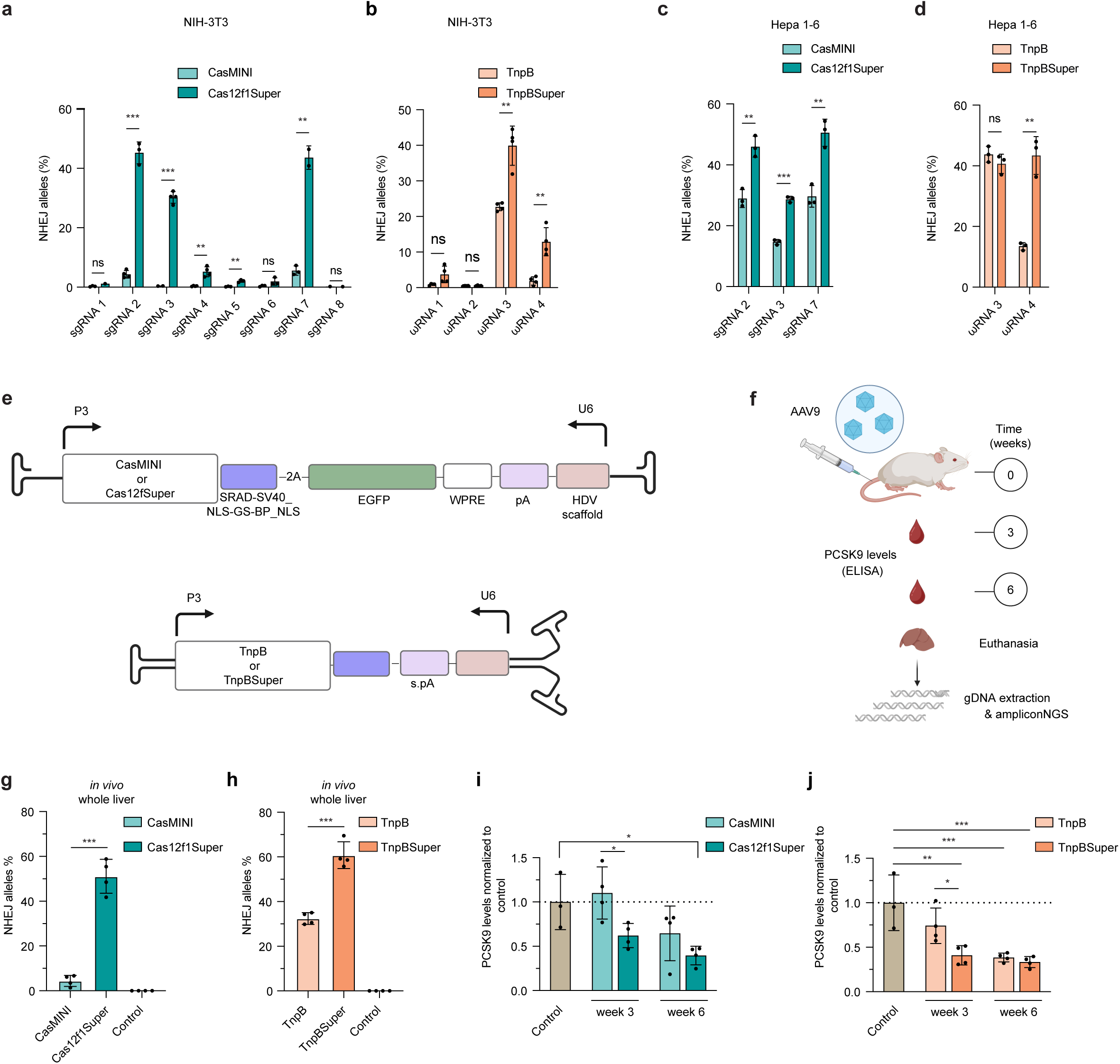
Efficient in vivo editing using AAV delivery of Cas12f1Super and TnpBSuper. (**a,b**) Absolute NHEJ efficiency in NIH-3T3 cells at targets within the *Pcsk9* gene upon editing with CasMINI/Cas12f1Super (**a**) or TnpB/TnpBSuper (**b**) measured with ampliconNGS (*n* = 3 biological replicates). (**c,d**) Absolute NHEJ efficiency in Hepa 1-6 cells at selected targets within the *Pcsk9* gene upon editing with CasMINI/Cas12f1Super (**c**) or TnpB/TnpBSuper measured with ampliconNGS (**d**) (*n* = 3 biological replicates). Each nuclease-guide RNA combination was delivered on a “all-in-one” plasmid. (**e**) Schematic representation of ssAAV9 vectors used for CasMINI/Cas12f1Super and scAAV9 vectors used for TnpB/TnpBSuper in *in vivo* experiments. Both designs are based on the TnpBmax architecture^19^. SRAD-SV40_NLS-GS-BP_NLS, linker from “TnpBmax” construct design; WPRE, woodchuck hepatitis virus post-transcriptional regulatory element; pA, polyadenylation signal sequence; s.pA, synthetic polyadenylation signal sequence. (**f**) Schematic overview of the *in vivo* experimental setup. AAV9 capsids were administered to mice via tail vein injection. PCSK9 protein levels were measured in blood serum at weeks 3 and 6. At week 6, murine livers were harvested and analyzed by amplicon ampliconNGS. (**g,h**) Editing efficiencies in whole liver at the 6-week timepoint following treatment with CasMINI/Cas12f1Super (**g**) or TnpB/TnpBSuper (**h**), measured by amplicon NGS (*n* = 4 animals). **(i,j)** PCSK9 protein levels following treatment with CasMINI/Cas12f1Super (**i**) or TnpB/TnpBSuper (**j**), measured in blood serum by ELISA normalized to untreated (control) mice (*n* = 3. Each dot represents an individual biological replicate or animal, and bars represent the mean ± standard deviation. All *P* values were calculated using an unpaired, two-sided *t*-test. ns, not significant (P ≥ 0.05), \**P*L<L0.05, \*\**P*L<L0.01, \*\*\**P*L<L0.001.

For mouse *in vivo* experiments, we selected site 7 for CasMINI/Cas12f1Super and site 4 for TnpBSuper. CasMINI/Cas12f1Super were delivered using a single-stranded AAV9 (ssAAV9) design, whereas TnpB/TnpBSuper were packaged in a self-complementary AAV9 (scAAV9) format, both based on previously optimized TnpBmax architectures^19^ (**Fig. 4e**).

AAV vectors were administered via tail vein injection to healthy C57BL/6J mice and circulating PCSK9 levels were quantified by ELISA at 3- and 6-week timepoints post-injection. Editing efficiency in the liver was assessed at the 6-week timepoint (**Fig. 4f**). For both systems, we observed significantly increased editing with Cas12f1Super and TnpBSuper compared to their respective wild-type counterparts (**Fig. 4g,h**). Editing efficiencies in whole liver reached 51±8% for Cas12f1Super and 61±6% for TnpBSuper, reflecting marked improvements over the parental constructs. This improved activity was reflected in more rapid drops in plasma PCSK9 protein levels for both Cas12f1Super and TnpBSuper, though after 6 weeks all systems approached a consistent minimum amount of PCSK9 (**Fig. 4i,j**).

Together, these results demonstrate efficient in vivo editing of a therapeutically relevant target by Cas12f1Super and TnpBSuper.

### Improved variant dCas12f1Super outperforms dCasMINI as adenine base editor

Encouraged by the high activity of our editing enzymes, we explored their potential applications in base editing. The additional enzymatic machinery appended to Cas proteins that enables base editing makes miniature editors particularly attractive in this context. Following the example of an existing dCasMINI-based adenine base editor^3^, we introduced two inactivating mutations (D326A, D510A) to generate dCasMINI and dCas12f1Super and fused TadA* from ABE8e to their N-termini^25^. We individually co-delivered these base editors to HEK293T cells as plasmids together with sgRNA-encoding plasmids targeting the genomically integrated Target CM1 or three endogenous loci (*TAOK1*, *P2RX5* and *TTR*) (**Fig. 5a**). We measured base editing using ampliconNGS. dCas12f1Super consistently outperformed dCasMINI fusion at all four sites, with up to a 10±2.5-fold increase at *TAOK1* (**Fig. S11a**) and reaching 17±0.1% absolute base editing at *TTR* (**Fig. 5b**). The per-base editing window remained similar between dCasMINI and dCas12f1Super, but with markedly increased activity for the variant (**Fig. 5c-f**). We observed almost exclusively A-to-G conversion, even at the most efficient *TTR* site (**Fig. 5g**). Notably, we observed zero indels even at the most efficient *TTR* target site, consistent with the use of catalytically inactive editor (**Fig. 5h, Fig. S11b**) This activity and product purity is encouraging for the potential use of Cas12f1Super beyond its evolved HDR and NHEJ context.

**Figure 5.**
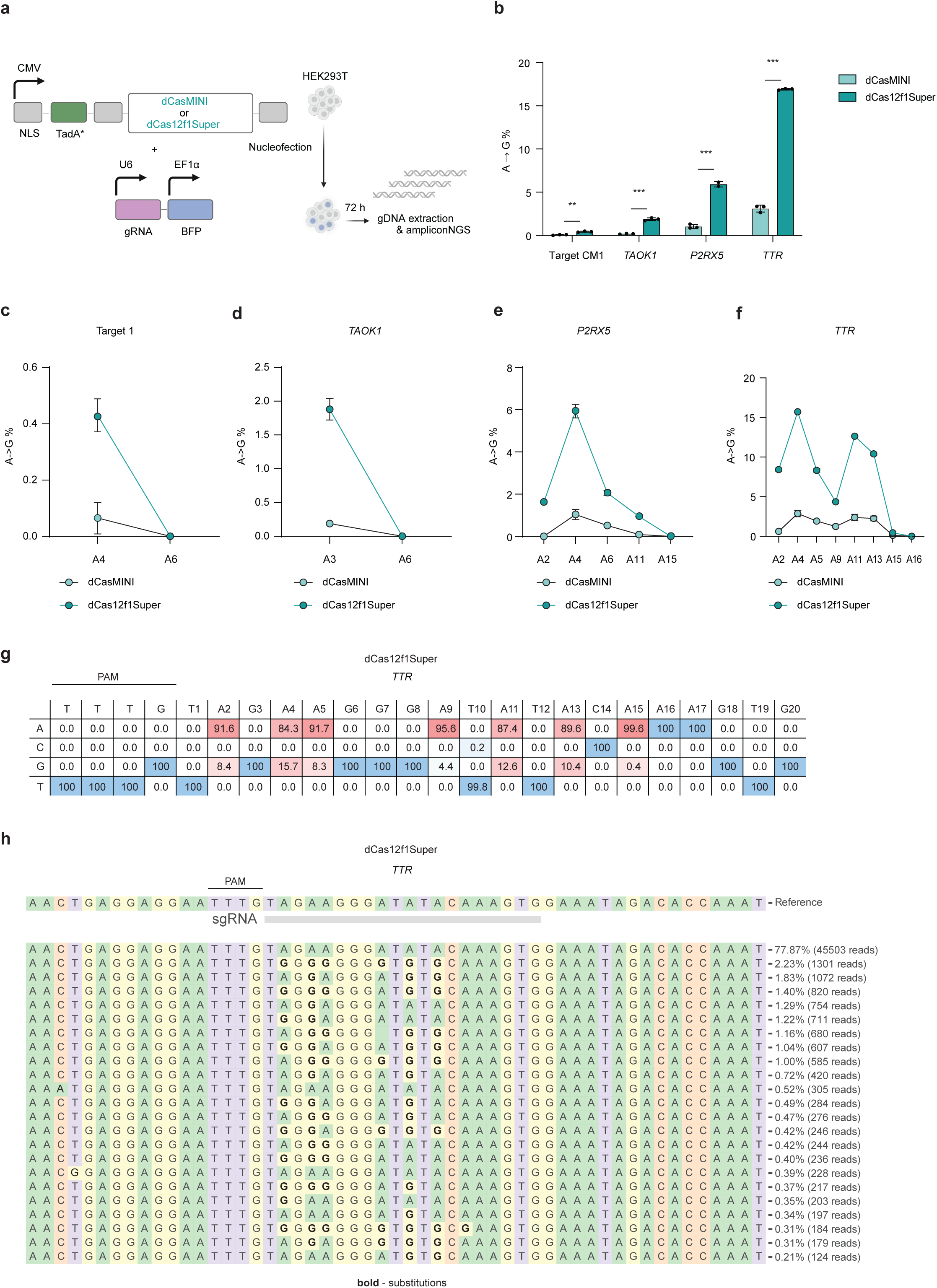
Improved variant dCas12f1Super outperforms dCasMINI as an adenine base editor. (**a**) Schematic showing adenine base editing of Target CM1 and endogenous sites in HEK293T using catalytically-dead CasMINI or Cas12f1Super (dCasMINI or dCas12f1Super, respectively). Plasmid expressing catalytically inactive enzyme fused with TadA* is co-delivered with guide-expressing plasmid. AmpliconNGS was performed 3 days post-transfection. (**b**) Overall adenine base editing activity of dCasMINI and dCas12f1Super at Target CM1 and indicated endogenous sites (*n* = 3 biological replicates). (**c–f**) Per-site adenine editing efficiencies of adenine base editors dCasMINI and dCas12f1Super at indicated endogenous genomic sites in HEK293T cells (*n* = 3 biological replicates). Editing outcomes were quantified as A•T-to-G•C conversion rates. (**g**) Product distributions at *TTR* endogenous site in HEK293T cells edited using a combination of adenine base editor dCas12f1Super and the corresponding sgRNA. (**h**) Allele plot at *TTR* endogenous site in HEK293T cells edited using a combination of adenine base editor dCas12f1Super and the corresponding sgRNA Each dot represents an individual biological replicate, and bars represent the mean ± standard deviation. All *P* values were calculated using an unpaired, two-sided *t*-test. ns, not significant (P ≥ 0.05), \**P*L<L0.05, \*\**P*L<L0.01, \*\*\**P*L<L0.001.

## Discussion

Progress in genome editing relies on the development of more active tools for mammalian (human) systems. While rational design allows direct testing in human cells, it is constrained by limited variant throughput and the challenge of predicting beneficial mutations from structure alone. Directed evolution in bacterial systems offers high throughput, but nucleases evolved in bacteria can underperform in human cells. Several studies have reported false positives, where mutations that appeared promising in bacterial cells failed to translate to human cells^26–28^. This may be due to differences in cellular context, including chromatin structure, DNA repair pathways and post-translational modifications.

These differences have motivated the development of eukaryotic platforms for directed evolution. Yeast-based systems partially address this gap^29^, but the biological differences between yeast and human cells still make functional divergence likely. DMS in human cells, such as a recent screen of AsCas12f1, illustrates the power of selection directly in the relevant context^8^. However, DMS in the context of human cell culture is only feasible for single point mutations and does not capture the potential of synergistic combinations that may emerge during selection. In this work, we developed a mammalian-based selection strategy to enrich for highly active multi-mutant variants and applied it to CasMINI and TnpB, yielding Cas12f1Super and TnpBSuper with strongly enhanced activity in human cells.

For Cas12f1Super, four out of five mutations involve a switch from positively to negatively charged residues. One example, K52E, lies near the sgRNA scaffold and improves activity despite losing a positive charge which is an unexpected result given the presumed importance of electrostatic interactions in that region. Given K52’s proximity to the sgRNA-interacting R66, its beneficial effect may result from stabilizing sgRNA binding by buttressing R66. The other charge-altering mutations could also reconfigure the protein to enhance nucleic acid engagement, though such effects are not readily apparent from the nuclease structure. E206K adds a positive charge near DNA, and a similar substitution (E206R) was previously proposed in a rational design study to enhance DNA binding^16^. In TnpBSuper, five of nine mutations originate from a single variant straight from selection (V322). Among the remaining mutations, E73K is positioned near DNA and may participate in protein-DNA interactions. Others include conservative changes such as L155V, L172V, V192L, and G299A. L172 is involved in stabilizing the TAM-proximal end of the heteroduplex^30^, suggesting that L172V may support more efficient duplex formation. While this manuscript was in revision, deep mutational scanning of ISDra2 TnpB individual point mutations in yeast identified some overlapping beneficial mutations (e.g. V192L) and some partially convergent mutations (e.g. L172G vs. L172V)^31^. However, the majority of mutations reported here are distinct from those identified by yeast-based selection. These differences likely reflect distinct selection environments (yeast vs. mammalian cell biology), non-saturating but combinatorial mutagenesis in our approach, and locus-dependent effects.

Our mammalian selection strategy focused on promoting HDR without directly suppressing NHEJ and the resulting variants showed higher HDR and NHEJ outcomes. Explicitly including HDR during selection and screening yielded TnpB and Cas12f1 variants with high overall activity, retaining the ability to perform both error-prone and precise editing. Cas12f1Super exhibited identical *in vitro* cleavage patterns and enzymatic parameters to CasMINI, and suggesting that its improved activity might stem from mammalian cell-specific factors such as increased expression, greater protein stability, or even interaction with the DNA repair machinery. Additionally, altered trans ssODN-cleavage activity of these nucleases could affect the availability of the donor template during repair. Nevertheless, these mechanisms remain speculative and require additional investigation.

In summary, our work demonstrates the power of mammalian-based selection to evolve compact, highly active genome editors tailored for function in complex cellular environments. The functional improvements observed in CasMINI and TnpB systems after their engineering in mammalian cells, including enhanced HDR, NHEJ, and base editing activity, together with their compatibility with AAV delivery, highlight their utility for broad applications in therapeutic genome engineering.

## Methods

### Cell culture

HEK293T, NIH-3T3, Hepa 1-6, and HEK293T reporter cell lines were cultured in Dulbecco’s Modified Eagle Medium (DMEM) with GlutaMAX (Thermo Fisher Scientific), supplemented with 10% fetal bovine serum (FBS, Thermo Fisher Scientific) and 100 U/mL penicillin-streptomycin (Gibco). HEK293T and NIH-3T3 cells were obtained from certified vendors, and Hepa 1-6 cells were provided by the laboratory of Prof. Dr. Markus Stoffel (ETH Zürich). These cell lines were maintained at 37°C in a humidified incubator with 5% CO[ and routinely tested for mycoplasma contamination using the MycoAlert kit (Lonza). Cells were never used beyond passage 20. Primary human T cells were isolated from Leukopaks (StemCell Technologies) using the EasySep™ Human T Cell Enrichment Kit, and activated by culturing in T25 flasks coated overnight at 4 °C with anti-CD3 (clone OKT3, functional grade; Thermo Fisher Scientific) and anti-CD28 (clone CD28.2, BioLegend) antibodies (1 µg/mL each in ultrapure H[O). Prior to electroporation, activated T cells were maintained for 48 hours in CTS medium (Advanced RPMI 1640 (44.5%), Click’s Medium (44.5%), FBS (10%), and 200 mM L-glutamine (1%)) supplemented with IL-7 (10 ng/mL) and IL-15 (5 ng/mL).

### HDR reporter cell line generation

Lentiviral packaging of HDR reporter constructs was performed in HEK293T cells using polyethyleneimine (PEI) as the transfection reagent. The HDR construct plasmid was co-transfected with the envelope plasmid (pCMV-VSV-G) and the packaging plasmid (pCMV-dR8.2 dvpr) at a 1:1:1 molar ratio. Viral supernatant was collected 48 hours post-transfection, filtered through a 0.45 µm filter, and used to transduce HEK293T cells with different HDR reporters. Reporter construct with Target CM1 was transduced with MOI of 6, while all other reporter constructs (Target CM2, Target TB1, Target TB2) were transduced with MOI of 0.2. Twenty-four hours after transduction media was replaced. Forty-eight hours after transduction, blasticidin selection was initiated at concentration of 6 µg/mL and continued for 3 weeks to establish stable cell lines.

### Cloning variant libraries

Libraries were generated by error-prone PCR (epPCR) using the GeneMorph II kit (Agilent) on CasMINI or TnpB-containing amplicons. Conditions were optimized to introduce ∼2 amino acid-changing mutations per open reading frame (ORF). PCR products and destination vectors were digested with AscI and BamHI for 4 hours, followed by overnight ligation at 16°C. Ligated products were isopropanol-precipitated and electroporated into MegaX DH10B cells (Thermo Fisher Scientific) using a Bio-Rad electroporator. After a 1-hour recovery in SOC medium, transformed cells were plated on large square agar plates and incubated overnight at 30°C. Colonies were harvested, and plasmid DNA was purified using a Qiagen Midiprep kit.

### Workflow for HDR-enriched variant selection

Variant libraries and corresponding wild-type (WT) plasmids were packaged into lentivirus using PEI-mediated co-transfection with the envelope plasmid (pCMV-VSV-G) and the packaging plasmid (pCMV-dR8.2 dvpr) in HEK293T cells at a 1:1:1 molar ratio. Lentiviral supernatants were used to transduce HEK293T HDR reporter cells. Forty-eight hours post-transduction, cells underwent two consecutive 2-day puromycin selection rounds (concentration 1.5 µg/ml). After 1 day of recovery, cells were electroporated with a guide RNA targeting the HDR reporter (Target CM1 for CasMINI or Target TB1 for TnpB) and an ssODN donor template to restore *EGFP* expression. Electroporations were performed using Lonza Amaxa 4D Nucleofector in 100 µL single Nucleocuvettes. Three days post-electroporation, EGFP-positive cells were isolated by FACS. Genomic DNA was extracted using the Gentra Puregene Cell Kit (Qiagen). Variants were PCR-amplified from genomic DNA, digested with AscI and BamHI, and cloned to generate the next library generation. The primers for amplifying variants from genomic DNA are listed in **Supplementary Table 2.**

### Design of ssODN donors

All ssODNs used for HDR at reporter and endogenous targets were 180 nucleotides long, complementary to the targeted strand, and symmetric around the cut site. For CasMINI or TnpB HDR reporter targets, the ssODNs included the 2-nucleotide substitution required to restore the *EGFP* start codon, along with an additional modification within the protospacer (positions 11–15 relative to the PAM or TAM sequence for CasMINI or TnpB, respectively). For CasMINI or TnpB endogenous targets, ssODNs were designed to include: (1) a 2-nucleotide substitution near the cut site (positions 23 & 24 relative to the PAM for CasMINI, or positions 17 & 18 relative to the TAM for TnpB); (2) a substitution within the protospacer (positions 11–15 for CasMINI, or 11–16 for TnpB); and (3) a PAM/TAM mutation (TTTR→TGTR for CasMINI, or TTGAT→TTGTT for TnpB). The protospacer and PAM/TAM modifications were included to prevent re-targeting following successful HDR. Final editing efficiency was quantified as the percentage of reads containing the ATG restoration (when editing HDR reporter cell lines) or the 2-nucleotide substitution near the cut site (when editing endogenous targets in HEK293T cell line). For SpCas9 endogenous targets, ssODNs were designed to include (1) a 2-nucleotide substitution near the cut site (positions 3 & 4 relative to the PAM); and (2) a PAM mutation (NGG→NTG).

ssODNs were ordered as Ultramer DNA oligos from Integrated DNA Technologies (IDT). For electroporating T cells, chemically protected ssODNs were used (ordered with four phosphorothioate modifications at the 5′ and 3′ ends). The sequences of ssODNs are listed in **Supplementary Table 3**.

### Testing workflow for HDR-enriched variant selection

The workflow for HDR-enriched variant selection was tested at Target CM1 with CasMINI. CasMINI was delivered via lentivirus, produced by PEI-mediated co-transfection of the envelope plasmid (pCMV-VSV-G), the packaging plasmid (pCMV-dR8.2 dvpr) and the CasMINI expression plasmid into HEK293T cells at a 1:1:1 molar ratio. The resulting lentiviral supernatant was used to transduce HEK293T HDR reporter cells (with Target CM1). Forty-eight hours post-transduction, cells underwent two consecutive rounds of 2-day puromycin selection (concentration 1.5 µg/ml). After 1 day of recovery, cells were electroporated with a guide RNA targeting the HDR reporter (Target CM1) and an ssODN donor template to restore *EGFP* expression. After 3 days cells were collected for flow cytometry analysis and genomic DNA extraction for subsequent ampliconNGS.

### Flow cytometry and fluorescence-activated cell sorting (FACS)

For all flow cytometry and FACS experiments cells were harvested 3 days post-transfection by dissociation with 0.05% Trypsin and resuspension in PBS supplemented with 10% FBS. Analytical flow cytometry for quantifying editing at HDR reporter targets in HEK293T-derived cell lines was performed using an Attune NxT Flow Cytometer (Thermo Fisher Scientific), and data were analyzed using FlowJo software (version 10.10.0). For all experiments in HEK293T cells at HDR reporter targets except for experiment involving TnpBmax in **Fig. S5f** cells were sequentially gated for live cells (FSC/SSC), singlets (FSC-A/FSC-H), BFP+, mCherry+, and finally EGFP+ populations **(Fig. S12a)**. For experiment involving comparison with TnpBmax in HEK293T cells in **Fig. S5f** cells were sequentially gated for live cells (FSC/SSC), singlets (FSC-A/FSC-H), mCherry+, and finally EGFP+ populations **(Fig. S12b)**. FACS for analysis of editing at endogenous sites in HEK293T, NIH-3T3 and Hepa 1-6 cell lines and FACS during big scale selections in HEK293T cells was performed using a BD FACSAria Fusion or BD FACSAria III. In case of FACS for analysis of editing at endogenous sites in HEK293T involving comparison with TnpBmax (**Fig. S5g)**, editing at TRAC locus (**Fig. S8d**) and comparison with SpCas9 (**Fig. 3j**, **S8g**) cells were sequentially gated for live (FSC/SSC), singlets (FSC-A/FSC-H), and finally mCherry+ populations **(Fig. S12b),** then 1 × 10^5^ cells were processed for genomic DNA extraction and subsequent ampliconNGS. In case of FACS for analysis of editing at endogenous sites in Hepa 1-6 cell line, cells were sequentially gated for live (FSC/SSC), singlets (FSC-A/FSC-H), and finally mCherry+ populations **(Fig. S12b),** then 1.2 × 10^4^ mCherry+ cells were sorted, cultured for 5 days, and 1 × 10^5^ cells were processed for genomic DNA extraction and subsequent ampliconNGS. In case of FACS for analysis of editing at endogenous sites in HEK293T (except for comparison with TnpBmax in **Fig. S5g,** editing at TRAC locus in **Fig. S8d** and comparison with SpCas9 in **Fig. 3j**, **S8g**) and in NIH-3T3 cell lines, cells were sequentially gated for live (FSC/SSC), singlets (FSC-A/FSC-H), and finally mCherry+/BFP+ population **(Fig. S13a)**, then 1 × 10^5^ BFP^+^/mCherry^+^ double positive cells were sorted and processed for genomic DNA extraction and subsequent ampliconNGS. In case of FACS during big scale selections in HEK293T cell line, cells were sequentially gated for live cells (FSC/SSC), singlets, SYTOX^TM^-Red-negative, and finally mCherry+/EGFP+ populations **(Fig. S13b)**, then all EGFP-positive cells were sorted from 1.1 × 10^8^ mCherry+ cells during round one and two of selection or from 2 × 10^7^ mCherry+ cells during rounds three and four of selection and processed for genomic DNA extraction.

### mRNA synthesis for T cell electroporation

mRNA encoding either TnpB or TnpBSuper was synthesized using the mMESSAGE mMACHINE™ T3 Transcription Kit (Thermo Fisher AM1348). The expression plasmid containing the T3 promoter was linearized, purified, and used for *in vitro* transcription. The resulting mRNA was purified with Monarch Spin RNA Cleanup Kit (New England Biolabs T2050L) and quantified by Qubit using the Qubit RNA Broad Range Kit (Thermo Fisher Q10210). The integrity of the mRNA constructs was verified with an automated electrophoresis system (Agilent).

### Electroporation of mammalian cells

To assess editing at endogenous sites in HEK293T, HEK293T-derived, and NIH-3T3 cell lines, 1-2[×[10[ cells were electroporated per reaction with a total of 600 ng plasmid DNA at a 1:1 molar ratio of nuclease-expression and guide RNA-expression plasmids, carrying mCherry and BFP markers, respectively. When the nuclease was stably expressed in HEK293T-derived lines, 200 ng of guide RNA-expressing plasmid (without a fluorescent marker) was used. 100[pmol of ssODN was co-electroporated when transfecting HEK293T and HEK293T-derived lines. For editing in activated primary T cells, 1[×[10^6^ cells were electroporated with 150[pmol of synthetic ωRNA (modified with 2’-O-methyl analogs on the first and last three bases and 3’ phosphorothioate internucleotide linkages between the first three and last two bases; Synthego), 4 µg of mRNA encoding the nuclease and 200 pmol of chemically protected ssODN. Electroporations were performed using the Lonza Amaxa 4D Nucleofector with 20 µL Nucleocuvette Strips. Cell lines were electroporated with SF buffer using program DG-130 (HEK293T and derivatives) or SG buffer using EN-158 (NIH-3T3). T cells were electroporated with P3 buffer using program EH-115. Following electroporation, HEK293T, HEK293T-derived, and NIH-3T3 cells were cultured for 3 days before flow cytometry or FACS. For T cells, the medium was exchanged 24 hours post-electroporation, and genomic DNA was harvested 3 days after the media change.

### Lipofection of mammalian cells

To assess editing at endogenous sites in the Hepa 1-6 cell line, 1.5 × 10[ cells were seeded per well in a 24-well plate and transfected with 1000 ng of an “all-in-one” plasmid expressing both the protein and guide RNA, along with an mCherry marker, using Lipofectamine 3000 (Thermo Fisher). Cells were cultured for 3 days prior to FACS.

### Base editing

Two inactivating mutations (D326A, D510A) were introduced to CasMINI or Cas12f1Super to get catalytically inactive variants: dCasMINI and dCas12f1Super. All sites in HEK293T cells were base-edited by co-electroporation of dCasMINI or dCas12f1Super-expressing plasmid and sgRNA-expressing plasmid as described above in the “electroporation of mammalian cells” section of these Methods. Cells were collected for ampliconNGS 3 days after transfection.

### Amplicon next-generation sequencing (ampliconNGS)

Three days post-transfection (4 days for T cells), cells were dissociated, resuspended in PBS + 10% FBS, and 1 × 10^5^ cells were pelleted and resuspended in 50 µL QuickExtract solution (Lucigen). Genomic DNA was extracted by incubation at 65°C for 30 min, followed by 98°C for 15 min. Amplicon libraries were prepared using a two-step PCR protocol. In PCR1, the target locus was amplified with primers containing Illumina adapter sequences. Products were validated on agarose gel and purified using SPRI beads (0.9× volume). In PCR2, indexing primers were added. Final libraries were bead-purified (0.7× volume), pooled, and sequenced using paired-end 2×150 bp reads on an Illumina platform. NGS data were analyzed using CRISPResso2 (v2.3.1)^32^. For analysis of base editing outcomes overall editing was calculated as a percentage of A→G conversion in all reads that had had at least 0.1% representation. The primers used for ampliconNGS are listed in **Supplementary Table 2**.

### Tagmentation of libraries

Amplicons corresponding to either the original sequences of CasMINI or TnpB (referred to as “Ctrl”), or variants from libraries collected before selection (termed “before_R1”) and after rounds 1, 2, or 4 of selection (“R1,” “R2,” and “R4,” respectively), were generated by PCR. Each of these amplicons (1 ng input) was used to prepare sequencing libraries using the Nextera XT DNA Library Preparation Kit (Illumina), which includes enzymatic fragmentation and adapter tagging. The resulting libraries were subjected to high-throughput sequencing on an Illumina platform.

### Tagmented library analysis

Quality of raw sequencing files was checked with fastqc v0.11.9^33^ with default arguments and reports were combined using multiqc v1.12^34^. Raw reads were aligned to the respective custom reference with bowtie2 v2.4.4^35^ and arguments -p 60 and -local. Samtools v1.16.1^36^ was used to convert the alignments to BAM format, sort by genomic location and generate indices. The callvariants.sh tool from BBMap v38.95^37^ was used to call variants from all alignments from the same reference in multisample mode with the arguments multisample = t, ploid = 1, clearfilters, trimq = 25, minqualitymax = 25 and minmapqmax = 5. Variants were further filtered using bcftools v1.15.1^38^ with argument -v snps to only include SNPs. Variants were annotated using ensemble-vep v112.0^39^ using its docker image with custom references and default arguments. The annotated SNPs were further processed with a custom script in R v4.3.1^40^. Only SNPs with an allele fraction greater than or equal to 0.003 after R4 and with a before_R1/R4 ratio greater than 3.77 were kept. Plots were generated with base R plotting functions or with ComplexHeatmap v2.18^41^.

### Western blot analysis

Briefly, HEK293T cells were electroporated with HA-tagged CasMINI/CasMINI_CV20 or TnpB/TnpB_CV2-expressing plasmid and sg/ωRNA-expressing plasmid targeting Target CM1 (for CasMINI/CasMINI_CV20) or Target TB1 (for TnpB/TnpB_CV2) in triplicate. Electroporations were carried out using the Lonza Amaxa 4D Nucleofector system with 20[µl Nucleocuvette Strips. After 3 days, cells were collected and lysed in RIPA buffer (50 mM Tris-HCl, 150 mM NaCl, 0.25% deoxycholic acid, 1% NP-40, 1 mM EDTA, pH 7.4) supplemented with protease inhibitors (Halt protease inhibitor cocktail, Thermo Fisher Scientific). Proteins were separated by polyacrylamide gel electrophoresis (PAGE) and transferred onto 0.2 µm nitrocellulose membranes using standard wet transfer methods. Membranes were blocked in TBS-T (150 mM NaCl, 20 mM Tris, 0.1% Tween-20, pH 7.4) containing 5% skim milk, then incubated with primary antibodies diluted in TBS-T supplemented with 5% BSA and 0.05% sodium azide. The primary antibodies used were anti-HA (Cell Signaling Technology, 3724) and anti-GAPDH (Cell Signaling Technology, 2118), both at 1:1,000 dilution. Detection was performed using fluorescent secondary antibody (LI-COR Biosciences, 926-32213) at 1:10,000 dilution. Blots were imaged using the Odyssey CLx scanner (LI-COR Biosciences).

### EVOLVEpro

The identities and relative activities of the best-performing single mutants of CasMINI (16 point mutants) and TnpB (7 point mutants) were used as input for EVOLVEpro (https://github.com/mat10d/EvolvePro)^17^. The top eight predicted activity-enhancing mutations for CasMINI and TnpB were individually cloned onto the Cas12f1Super and TnpBSuper backgrounds. These new variants were tested in HEK293T cells at Target CM1 and Target CM2 (for CasMINI variants) or Target TB1 and Target TB2 (for TnpB variants) via co-delivery of a nuclease-expressing plasmid, a sg/ωRNA-expressing plasmid and ssODN.

### DISCOVER-Seq

HEK293T cells (2 × 10^7^) were transfected with 120 µg of each “all-in-one” construct using Lipofectamine 3000 (Thermo Fisher). Fifteen hours post-transfection, cells were pelleted and resuspended in room temperature RPMI (without supplements). Cells were crosslinked with 1% formaldehyde (Thermo Fisher) and incubated for 15 min at room temperature. Formaldehyde was quenched with 125 mM glycine for 3 min on ice. Cells were then pelleted at 4°C, washed twice with ice-cold PBS, and snap frozen in liquid nitrogen. Pellets were stored at -80°C prior to processing. MRE11 ChIP-Seq was performed next as described in Wienert et al^42^. Briefly, samples were thawed on ice and lysed using ice-cold buffers LB1, LB2, and LB3. The isolated DNA was sonicated to obtain ∼300 bp chromatin fragments using a Covaris S2 with the following settings: 12 cycles of duty cycle 5 %, intensity 5, 200 cycles per burst for 60 s. For each chromatin immunoprecipitation (ChIP), 10 μl of MRE11 antibody (PA3-16527R; Thermo Fisher) per ChIP was prebound to protein A Dynabeads (Invitrogen). MRE11-associated chromatin was immunoprecipitated with antibody-bound beads, rotating overnight at 4°C. Dynabeads were washed with RIPA buffer and the DNA was eluted by incubating overnight at 65°C in elution buffer containing SDS. For the final clean-up, the samples were digested with Proteinase K and RNase A in TE buffer for 1 h at 55°C. DNA fragments were purified using the MinElute Kit (Qiagen). Sequencing libraries were prepared using the NEBNext Ultra II workflow (New England BioLabs). Indexed libraries were pooled and sequenced on a NovaSeqX (Illumina) with 2×150 paired-end reads and a target depth of 40 million reads per sample. To identify genome editing target sites, NGS reads were first trimmed for the Illumina adapter sequences using cutadapt^43^ (v4.6), followed by alignment to the human genome (hg38) using bowtie2^44^ (v2.5.2). Resulting alignments were then deduplicated and subsampled to 40 million reads per sample using samtools^38^ (v1.19). Off-target site calling was performed using BLENDER^21,42^ (https://github.com/cornlab/blender) with custom parameters followed by manual inspection.

Plasmids used for DISCOVER-Seq are listed in **Supplementary Table 4.**

### CasMINI and Cas12f1Super purification

Both nucleases (His^6^-GFP-2xNLS-CasMINI/Cas12f1Super) were expressed from plasmids listed in **Supplementary Table 4**. Plasmids were transformed into *Escherichia coli* BL21 (DE3) cells, and protein expression was induced at an optical density (OD[[[) of 0.5 by adding IPTG to a final concentration of 0.5 mM. Cultures were then incubated at 18°C for 18 h. Cells were harvested and resuspended in buffer A (25 mM Tris-HCl (pH 7.6), 1 M NaCl, 5 mM β-mercaptoethanol) supplemented with 5% glycerol, 1 mM PMSF, 1 µg/ml pepstatin A, and 1 µg/ml leupeptin. Lysis was performed by sonication, and the lysates were clarified by centrifugation at 40,000 × *g* for 30 min. The supernatant was applied to Ni-NTA Superflow resin (Qiagen) pre-equilibrated with buffer A without supplements, washed with buffer A containing 25 mM imidazole, and eluted with buffer A containing 250 mM imidazole. The His^6^-GFP tag was removed by overnight digestion with TEV protease at 4°C. The resulting protein solution was diluted to reduce NaCl concentration to 0.2 M and loaded onto a HiTrap SP HP column (Cytiva) for ion exchange chromatography, with elution performed using a linear NaCl gradient (0.2–2 M). Finally, the protein fractions containing 2xNLS-CasMINI/Cas12f1Super were concentrated in a 30kDa molecular weight cut-off Amicon Ultra-15 Centrifugal Filter Unit (Merck) and further purified by size exclusion chromatography using a Superdex 200 column (GE Healthcare) pre-equilibrated with 25 mM Tris (pH 7.6), 200 mM NaCl, 2 mM DTT, 1 mM MgCl_2_. Fractions containing the target protein were pooled, diluted to a final concentration of 0.24 mg/ml, and stored at –80°C.

### sgRNA synthesis for plasmid DNA cleavage assay

The single guide RNA (sgRNA) templates were generated by two rounds of PCR using overlapping oligonucleotides containing the T7 promoter, sgRNA scaffold, and 20 nt targeting sequence. The obtained templates were then used for *in vitro* transcription (IVT) using the TranscriptAid T7 High Yield Transcription Kit (Thermo Fisher Scientific). The IVT reaction products were purified using the GeneJET RNA Purification Kit (Thermo Fisher Scientific). Oligonucleotides used for the template generation and sequences of the sgRNA templates, are listed in **Supplementary Tables 2** and **5, respectively**.

### CasMINI/Cas12f1Super-sgRNA complex assembly

CasMINI and Cas12f1Super ribonucleoprotein (RNP) complexes were assembled by mixing CasMINI or Cas12f1Super protein with sgRNA at 2:1 molar ratio and incubating at 37°C for 30 min in 10 mM Tris–HCl (pH 7.5 at 37°C), 100 mM NaCl, 1 mM DTT, and 1 mM EDTA buffer.

### Plasmid DNA cleavage assay

Plasmid DNA cleavage reactions were performed by mixing 100 nM of CasMINI-sgRNA complex with 3 nM of plasmid DNA (pTK1910 or pTK1911) in 2.5 mM Tris-HCl (pH 7.5 at 37°C), 25 mM NaCl, 10 mM MgCl_2_, 0.25 mM DTT, and 0.25 mM EDTA reaction buffer. The samples were incubated at 37-55°C, and the aliquots, removed at timed intervals (0, 1, 2, 5, 10 15, and 30 min), were mixed with 3×loading dye solution containing 75 mM EDTA, 0.1% SDS, and 0.03% Bromophenol Blue in 50% (v/v) glycerol. The samples were analyzed by running agarose gel electrophoresis and quantified using GelAnalyzer (v23.1.1). The linear cleavage products from the 30-minute endpoint aliquots were purified and subjected to run-off Sanger sequencing to determine the cleavage patterns on both the target and non-target DNA strands.

### Adeno-associated virus production

AAV9 vectors were produced by the Viral Vector Facility at the Neuroscience Center Zurich. In brief, vectors were purified by ultracentrifugation followed by diafiltration. Physical titers (vg[ml[¹) were quantified using a Qubit 3.0 fluorometer (Invitrogen). CasMINI/Cas12f1Super were packaged in a single-stranded AAV9 (ssAAV9) format, whereas TnpB/TnpBSuper constructs were packaged in a self-complementary AAV9 (scAAV9) format.

### Mouse studies

Animal experiments were performed following protocols approved by the Kantonales Veterinäramt Zürich and in compliance with the Swiss Animal Welfare Act and Protection ordinance. Mice (C57BL/6J) were housed in a pathogen-free animal facility at the Institute of Pharmacology and Toxicology at the University of Zurich and kept in a temperature- and humidity-controlled room (21[°C, 50% RH) on a 12-hour light-dark cycle and fed a standard laboratory chow (Kliba Nafag no. 3437 with 18.5% crude protein). AAV vectors were administered via tail vein injection in 6-week-old mice at a dose of 5 × 10^13^ vg/kg (n=4 per group, 2 males and 2 females). Mice were fasted for 3-4 hours before blood was collected from the tail vein for the 3-week timepoint or the inferior vena cava after the endpoint of the experiments.

### Serum biochemistry

Mouse PCSK9 protein levels in mouse plasma collected at 3 weeks and 6 weeks (endpoint) post-injection were determined using Mouse Proprotein Convertase 9/PCSK9 Quantikine ELISA Kit (R&D Systems MPC900) according to the instructions provided by the manufacturer. PCSK9 protein levels were normalized to those of untreated control mice. For each control mouse the 3- and 6-week timepoints were first averaged, and the resulting per-mouse values were then averaged across control mice to obtain a single normalization reference. One control mouse was excluded due to highly variable measurements between timepoints.

### Plasmid cloning and mutagenesis

#### Lentiviral reporter constructs

EFα-TAM/PAM-protospacer-EGFP-PGK-BSD fragment was inserted in a lentiviral backbone (Addgene ID 60954) using PCR and Gibson assembly. Protospacer is either Target CM1 or Target CM2 (for CasMINI reporter constructs) or Target TB1 or Target TB2 (for TnpB reporter constructs). *EGFP* gene contained a different triplet instead of a start codon: GGG for CasMINI reporters, AGA for TnpB reporters. Sequences or Target CM1, Target CM2, Target TB1, Target TB2 are provided in **Supplementary Table 2**.

#### Lentiviral CasMINI/TnpB expressing constructs

EF1α-2xNLS-CasMINI/TnpB-PGK-Puro-T2A-mCherry fragment was inserted in a lentiviral backbone (Addgene ID 52962) using PCR and Gibson assembly. These constructs were used as templates for epPCR for generating libraries of variants of CasMINI and TnpB. Mutations for single mutants (SMs) and combinatorial variants (CVs) were cloned using PCR mutagenesis and Gibson assembly.

#### sgRNA/ωRNA expressing constructs

Scaffolds of CasMINI or TnpB were cloned downstream from the human U6 promoter in a pUC19 plasmid. ge4.1 scaffold (100 nt) was used for CasMINI. Either the original (230 nt) or Trim2 scaffold (99 nt) was used for TnpB. The targeting part of the guide RNA sequence was cloned downstream from the scaffold sequence. EF1α-BFP fragment was cloned downstream from the guide. A version of these constructs without the EF1α-BFP part was used during HDR selection. Sequences of guide RNA scaffolds are provided in **Supplementary Table 6.**

“All-in-one” construct for transient co-expression of TnpB and Trim2 ωRNA (also called ωRNA*) was obtained from Addgene (ID 212967). “TnpBmax” design was cloned in in this backbone using Gibson assembly. This backbone was modified to co-express CasMINI and its sgRNA using Gibson cloning. “All-in-one” construct for transient co-expression of SpCas9 and its sgRNA was obtained from Addgene (ID 48138).

#### AAV constructs

The ssAAV construct used for *in vivo* experiments was kindly provided by the laboratory of Gerald Schwank. CasMINI or Cas12f1Super, together with the sgRNA, were cloned into a “TnpBmax”^19^ design using Gibson assembly.

The scAAV construct was obtained from the Viral Vector Facility of the University of Zurich (UZH), Zurich. TnpB or TnpBSuper, along with the ωRNA, were cloned into a “TnpBmax”^19^ design using Gibson assembly.

#### mRNA expressing constructs

NLS-TnpB/TnpBSuper-NLS fragment was placed under T3 promoter for subsequent *in vitro* transcription.

#### Base editor constructs for CasMINI

Two inactivating mutations (D326A, D510A) were introduced in CasMINI and Cas12f1Super with PCR to get dCasMINI and dCas12f1Super. Next, in ABE8e construct (Addgene ID 138489) dCas9 was replaced either with dCasMINI or dCas12f1Super.

#### Plasmid DNA substrates for cleavage assays

The plasmid DNA substrates (pTK1910 and pTK1911) for *in vitro* cleavage reactions were obtained by cloning PAM and target containing oligoduplex into a pUC18-based plasmid (pTK1909) through BpiI restriction endonuclease (Thermo Fisher Scientific) sites. The sequences of oligonucleotides are available in **Supplementary Table 2**.

For all Gibson assemblies, the New England Biolabs (NEB) HiFi Assembly Master Mix was used. All PCR reactions were performed using Q5 DNA Polymerase (NEB).

All plasmids used in this study are listed in **Supplementary Table 4.**

### Statistical analysis

GraphPad Prism 10.5.0 software was used for statistical analysis. Results are presented as mean ± standard deviation. All *P* values were calculated using an unpaired, two-sided *t*-test. Holm–Šidák correction was used for multiple comparisons. Schemes were created with Biorender.com. Protein structures were visualized with ChimeraX^45^.

## Data availability

Sequencing files for tagmented libraries and DISCOVER-Seq data are available in Sequence Read Archive BioProject PRJNA1286789.

## Acknowledgements

We thank Dr. Florian Mair, Dr. Malgorzata Kisielow, Anette Schütz, Renan Antonialli, Dr. Irini Vgenopoulou of the Flow Cytometry Core Facility of the ETH Zurich for cell sorting and support with flow cytometric analysis; Dr. Susanne Kreutzer of the ETH Zürich Genome Engineering and Measurement Lab for performing next-generation sequencing; Dr. M. Muhar, M. Schlapansky, Dr. Z. Kontarakis, Dr. E. Karasu and all of the other members or the Corn laboratory for helpful discussions; Prof. Dr. Markus Stoffel and Svenja Godbersen for gifting Hepa 1-6 cell line; Dr. Kim Marquart and the Viral Vector Facility of UZH (Zurich) for providing AAV construct backbones. J.E.C. is supported by the NOMIS Foundation and the Lotte und Adolf Hotz-Sprenger Stiftung. funded by the NOMIS Foundation and by the Lotte and Adolf Hotz-Sprenger Stiftung. T.K. is supported by “Mission-driven Implementation of Science and Innovation Programmes” No. 02-002-P-0001, funded by the Economic Revitalization and Resilience Enhancement Plan “New Generation Lithuania”. This project has received funding from the European Research Council (ERC) under the European Union’s Horizon 2020 research and innovation funding programmes (grant agreement no. 855741-DDREAMM-ERC-2019-SyG; no. 101057659; no. 101070740; no. 101057438), Swiss State Secretariat for Education, Research and Innovation (SERI), SNSF Project Funding grants (310030_188858 and 320030-227979) and SNF Bridge-Discovery grant (40B2-0_226603). LV.V. is supported by the ETH Pioneer Fellowship.

## Contributions

F.G. and J.E.C conceptualized the study. F.G. and I.S. performed most of the experiments. Y.-Y.L. purified the recombinant proteins. LV.V. and C.DY. performed DISCOVER-Seq and produced graphs in **Fig 3k,l**. A.T. performed the *in vivo* experiments. M.S.S. performed bioinformatic analyses and produced graphs in **Fig. 1d**. T.K. and G.D. performed *in vitro* biochemical studies and produced graphs in **Fig. S7**. L.V.B. and I.V. performed editing of primary T cells. F.G., A.G. and J.E.C. wrote the article. G.S. and V.S. reviewed and edited the article. J.E.C. supervised the study.

## Declaration of interests

J.E.C. is a co-founder of Serac Biosciences and a scientific advisory board (SAB) member of Relation Therapeutics and Hornet Bio. The laboratory of J.E.C. has had funded collaborations with Allogene and Cimeio. J.E.C. and F.G. are inventors on a patent filed by the ETH related to this work. The remaining authors declare no competing interests.

## Supplementary figure legends

**Figure S1.**
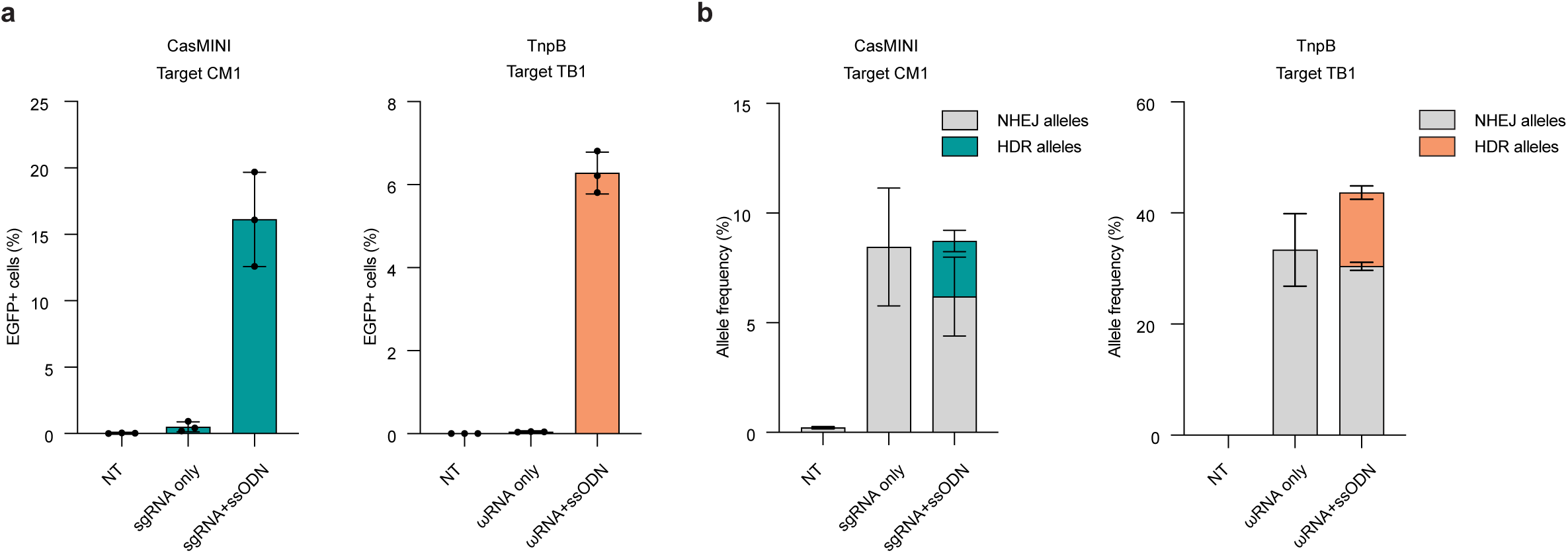
HDR reporter validation. (**a, b**) Testing of CasMINI and TnpB genome editors at two targets (Target CM1 for CasMINI and Target TB1 for TnpB) within HDR reporter’s framework in HEK293T cells (*n* = 3 biological replicates). EGFP percentages assessed by flow cytometry (**a**) and HDR and NHEJ allele percentages obtained through ampliconNGS (**b**) were used to quantify editing outcomes for indicated nucleases. NT, non-targeting. Each dot represents an individual biological replicate, and bars represent the mean ± standard deviation.

**Figure S2.**
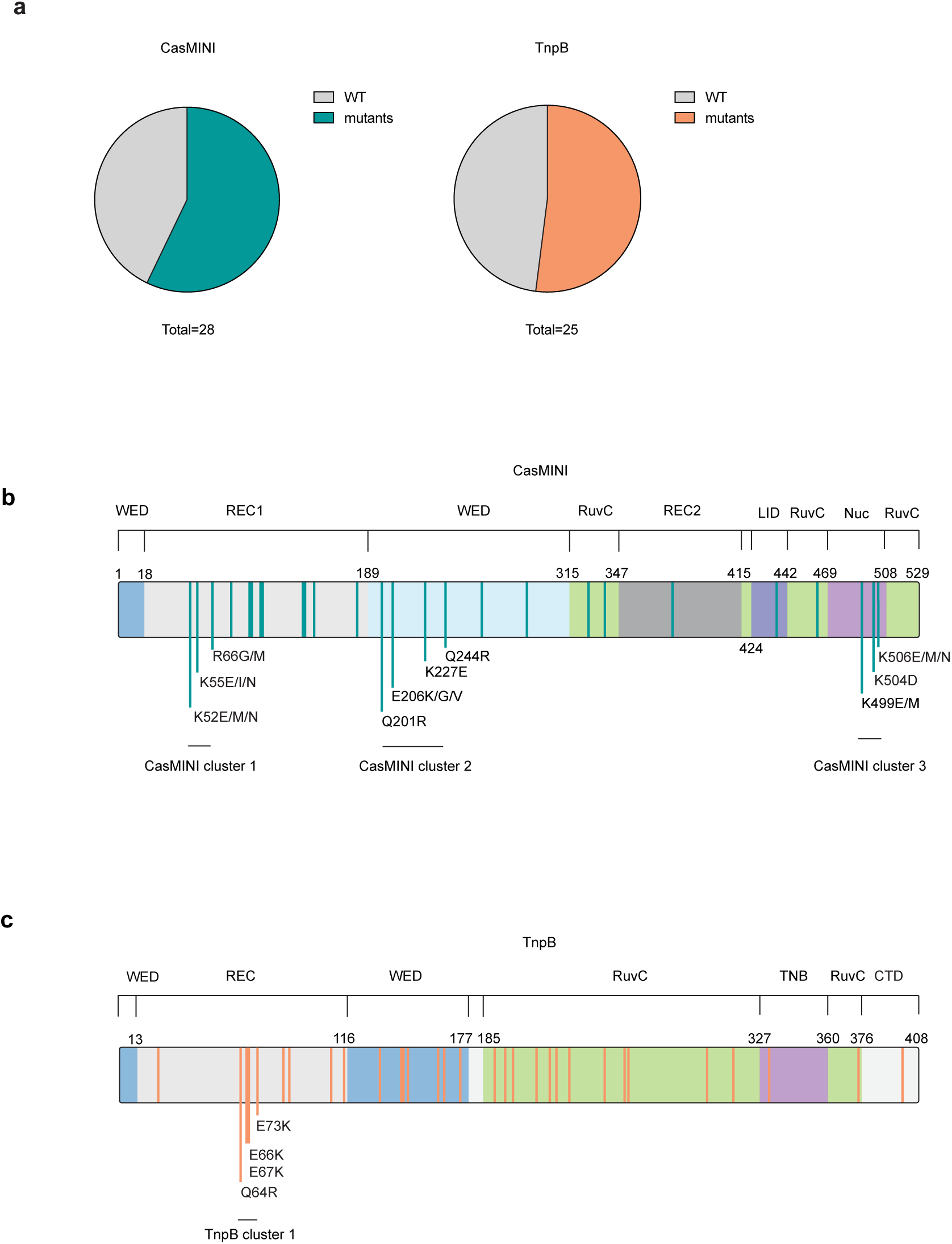
Localization of mutations enriched after selection for HDR with CasMINI and TnpB. (**a**) Pie chart showing proportion of unmutated (WT) and mutated (variants) sequences in libraries used for the first round of selection for indicated nucleases. For purpose of estimating this proportion 28 (for CasMINI) or 25 (for TnpB) randomly picked colonies from respective libraries were picked and analyzed by Sanger sequencing. (**b,c**) Mutations from Fig. 1d mapped on the domain organization of CasMINI (PDB: 7l49) (**b**) and TnpB (PDB: 8H1J) **(c).** The mutations from clusters of enriched mutations after round 4 of selection are labelled for each nuclease.

**Figure S3.**
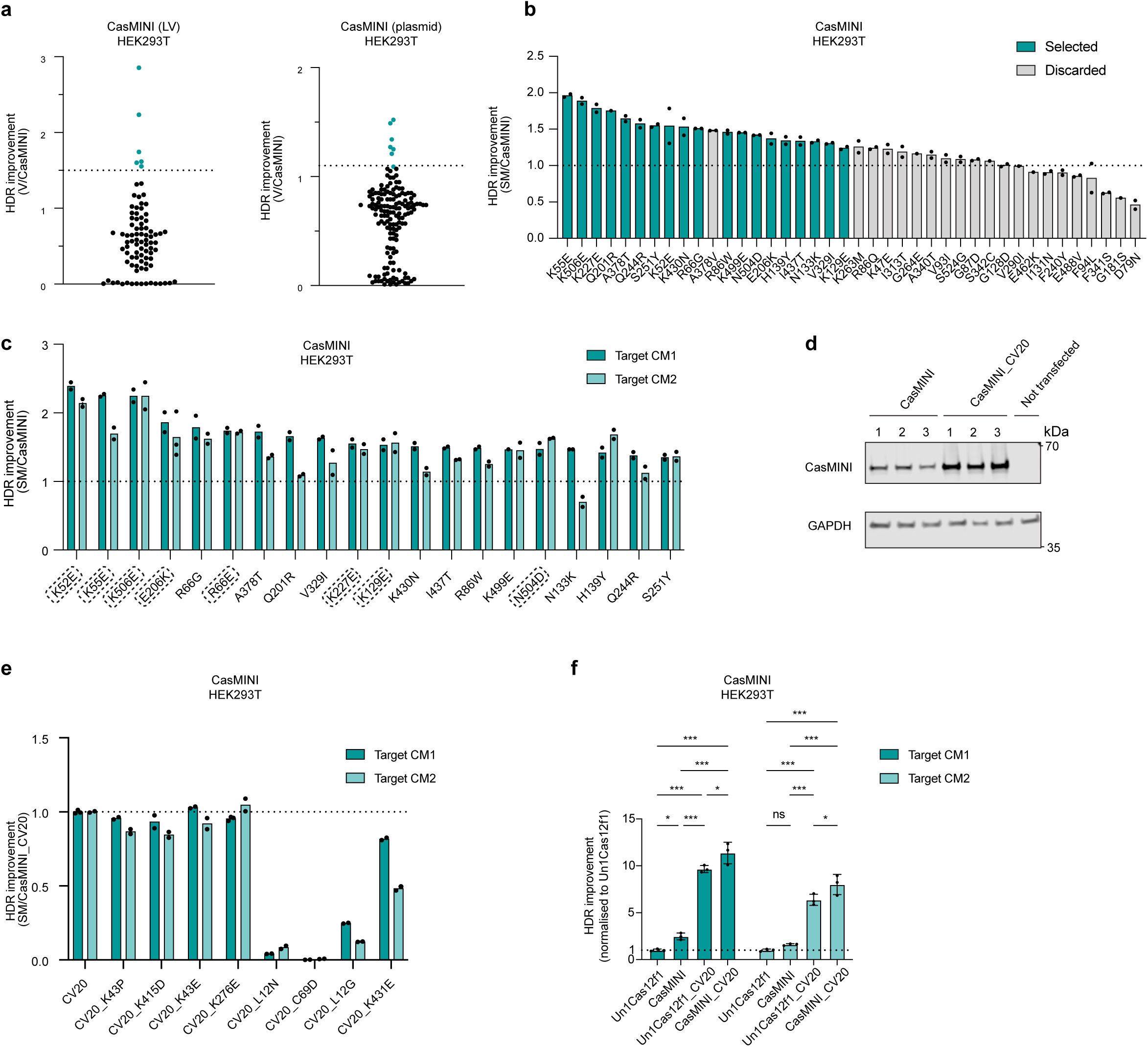
Selection of CasMINI mutations for generating combinatorial variants (CVs). (**a**) HDR efficiencies of 88 CasMINI variants were tested in a lentiviral (LV) format and 158 CasMINI variants tested with plasmid delivery at Target CM1 in HEK293T cells. HDR values were measured as percentage of EGFP-positive cells from BFP/mCherry double positive cells (BFP marks the guide-expressing construct; mCherry marks the nuclease-expressing construct) and normalized to CasMINI. Highlighted variants (teal) were selected for further validation using plasmid delivery in duplicate. (**b**) HDR efficiencies of single mutants (SMs) derived from selected high-performing CasMINI variants or enriched in NGS after round four of selection, tested at Target CM1 in HEK293T cells. HDR values were measured as percentage of EGFP-positive cells from BFP/mCherry double positive cells (BFP marks the guide-expressing construct; mCherry marks the nuclease-expressing construct) and normalized to CasMINI (*n* = 2 biological replicates). Highlighted SMs (teal) were selected for further testing at the orthogonal HDR reporter target (Target CM2). A378V was not selected due to the presence of A378T with similar or slightly higher editing efficiency, which was chosen for further analysis. Each dot represents an individual biological replicate, and bars represent the mean. (**c**) HDR efficiencies of selected CasMINI SMs tested in parallel at two distinct targets (Target CM1 and Target CM2) in HEK293T cells, measured as percentage of EGFP-positive cells from BFP/mCherry double positive cells (BFP marks the guide-expressing construct; mCherry marks the nuclease-expressing construct), with values normalized to CasMINI (*n* = 2 biological replicates). The mutations selected for making combinatorial variants are highlighted. Each dot represents an individual biological replicate, and bars represent the mean. (**d**) Western blot analysis of CasMINI and CasMINI_CV20 protein expression levels. Expression of each nuclease was performed in triplicate together with sgRNA for Target CM1 in HEK293T cells. GAPDH was used as a loading control. (**e**) HDR efficiencies of eight newly generated CasMINI variants, each containing one mutation derived from EVOLVEpro, cloned onto the CasMINI_CV20 background, tested in HEK293T cells (*n* = 2 biological replicates). HDR values were measured as percentage of EGFP-positive cells from BFP/mCherry double positive cells (BFP marks the guide-expressing construct; mCherry marks the nuclease-expressing construct) and normalized to the previously best-performing variant, CasMINI_CV20. Each dot represents an individual biological replicate, and bars represent the mean. (**f**) Comparison of editing efficiencies of Un1Cas12f1, CasMINI, Un1Cas12f1_CV20, CasMINI_CV20 at Target CM1 and Target CM2 in HEK293T cells. HDR values were measured as percentage of EGFP-positive cells from BFP/mCherry double positive cells (BFP marks the guide-expressing construct; mCherry marks the nuclease-expressing construct) and normalized to Un1Cas12f1 (*n* = 3 biological replicates). Each dot represents an individual biological replicate, and bars represent the mean ± standard deviation. All *P* values were calculated using an unpaired, two-sided *t*-test. ns, not significant (P ≥ 0.05), \**P*L<L0.05, \*\**P*L<L0.01, \*\*\**P*L<L0.001.

**Figure S4.**
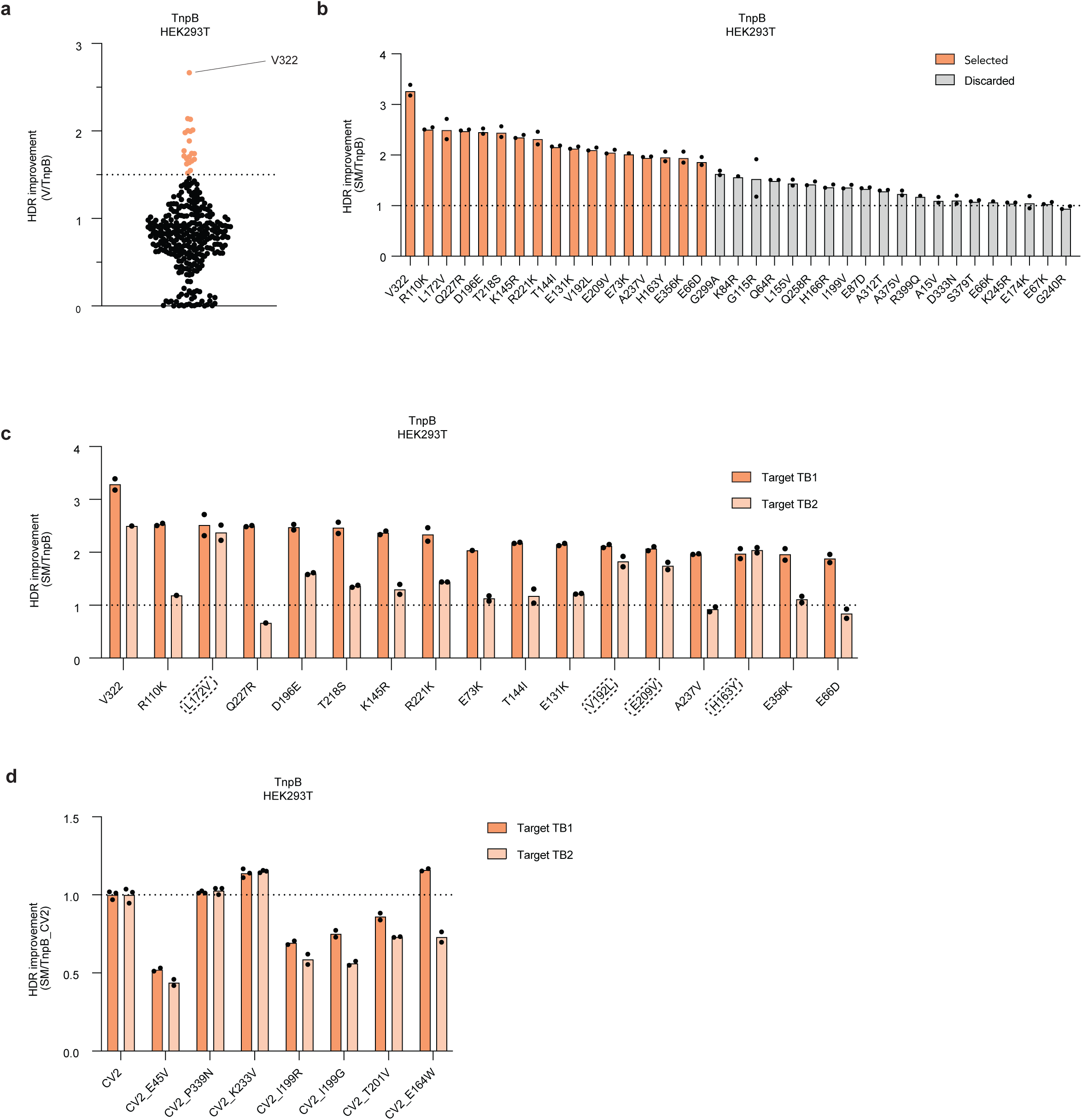
Selection of TnpB mutations for generating combinatorial variants (CVs). (**a**) HDR efficiencies of 311 TnpB variants with plasmid delivery at Target TB1 in HEK293T cells. HDR values were measured as a percentage of EGFP-positive cells from BFP/mCherry double positive cells (BFP marks the guide-expressing construct; mCherry marks the nuclease-expressing construct) and normalized to TnpB. Highlighted variants (orange) were selected for further validation using plasmid delivery in duplicate. Top-performing is indicated (V322). (**b**) HDR efficiencies of all TnpB single mutants (SMs) derived from selected high-performing TnpB variants or enriched in NGS after round four of selection, tested at Target TB1 in HEK293T cells (*n* = 2 biological replicates). HDR values were measured as percentage of EGFP-positive cells from BFP/mCherry double positive cells (BFP marks the guide-expressing construct; mCherry marks the nuclease-expressing construct) and normalized to TnpB. Highlighted SMs were selected for further testing at the orthogonal HDR reporter target (Target TB2). Each dot represents an individual biological replicate, and bars represent the mean. (**c**) HDR efficiencies of selected TnpB SMs tested in parallel at two distinct targets (Target TB1 and Target TB2) in HEK293T cells measured as percentage of EGFP-positive cells from BFP/mCherry double positive cells (BFP marks the guide-expressing construct; mCherry marks the nuclease-expressing construct), with values normalized to TnpB (*n* = 2 biological replicates). The mutations selected for making combinatorial variants are highlighted. Each dot represents an individual biological replicate, and bars represent the mean. (**d**) HDR efficiencies of seven newly generated TnpB variants, each containing one mutation derived from EVOLVEpro, cloned onto the TnpB_CV2 background, tested in HEK293T cells (*n* = 2 biological replicates). HDR values were measured as percentage of EGFP-positive cells from BFP/mCherry double positive cells (BFP marks the guide-expressing construct; mCherry marks the nuclease-expressing construct) and normalized to the previously best-performing variant, TnpB_CV2. Each dot represents an individual biological replicate, and bars represent the mean.

**Figure S5.**
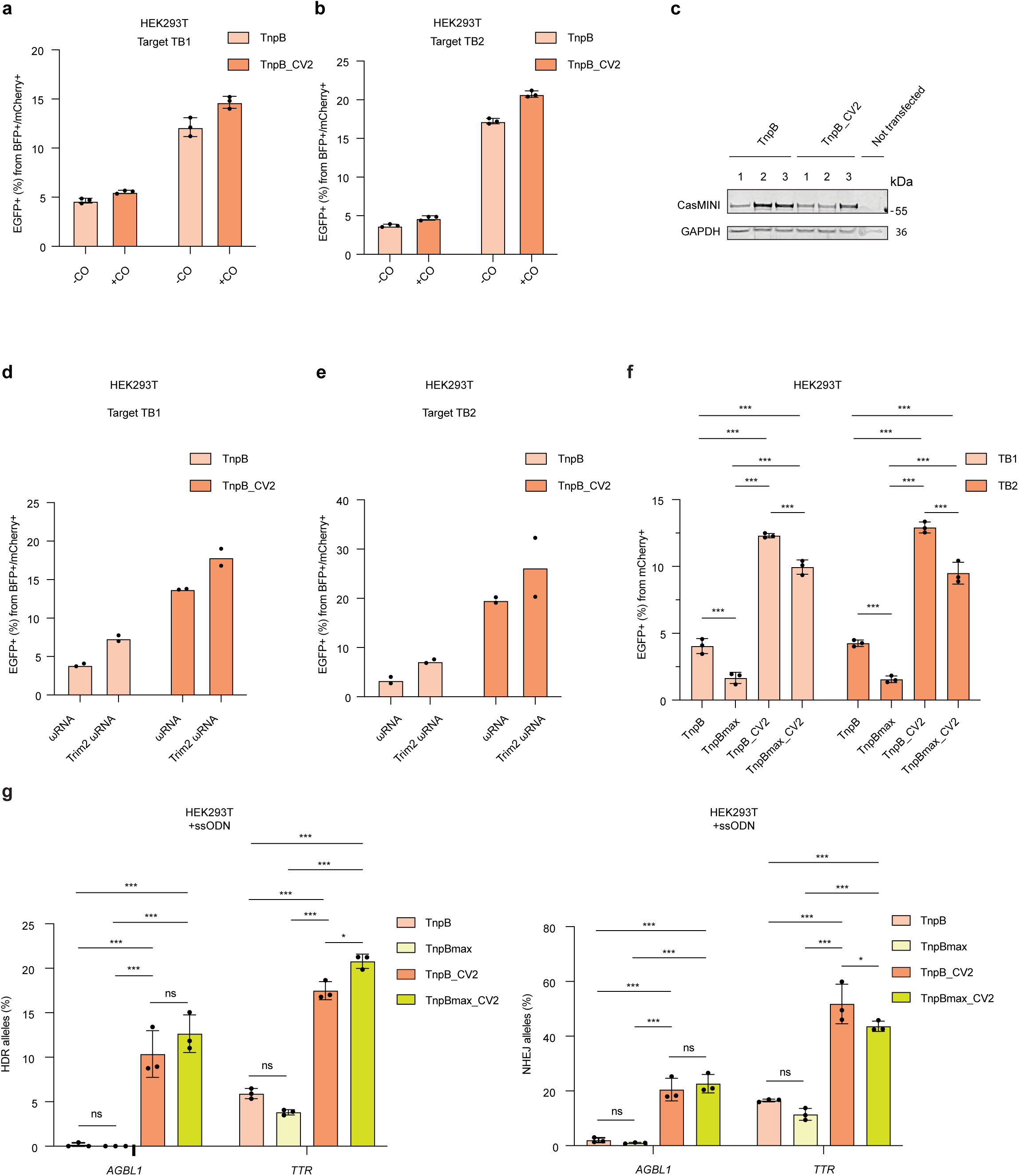
Mammalian codon-optimization and usage of shorter scaffold for TnpB enhance editing at HDR reporter targets in HEK293T cells. (**a,b**) HDR efficiencies with mammalian codon-optimized (+CO) and not codon-optimized (-CO) versions of TnpB and TnpB_CV2 in HEK293T cells at HDR reporter Target TB1 (**a**) and Target TB2 (**b**) (*n* = 3 biological replicates). HDR was measured as percentage of EGFP-positive cells from BFP/mCherry double positive cells (BFP marks the guide-expressing construct; mCherry marks the nuclease-expressing construct). Each dot represents an individual biological replicate, and bars represent the mean ± standard deviation. (**c**) Western blot analysis of mammalian codon-optimized TnpB and TnpB_CV2 protein expression levels (*n* = 3 biological replicates). Expression of each nuclease was performed together with ωRNA for Target TB1 in HEK293T cells. GAPDH was used as a loading control. (**d,e**) HDR efficiencies of mammalian codon-optimized TnpB and TnpB_CV2 combined with either Trim2 ωRNA scaffold or original ωRNA in HEK293T cells at HDR reporter Target TB1 (**d**) and Target TB2 (**e**). HDR was measured as percentage of EGFP-positive cells from BFP/mCherry double positive cells (BFP marks the guide-expressing construct; mCherry marks the nuclease-expressing construct) (*n* = 2 biological replicates). Each dot represents an individual biological replicate, and bars represent the mean. (**f**) HDR efficiencies of mammalian codon-optimized TnpB, TnpBmax, TnpB_CV2 and TnpBmax_CV2 combined with Trim2 ωRNA scaffold in HEK293T cells at HDR reporter targets TB1 and TB2. HDR was measured as percentage of EGFP-positive cells from mCherry+ cells (all-in-one construct was used for this experiment) (*n* = 3 biological replicates). (**g**) HDR and NHEJ efficiencies of mammalian codon-optimized TnpB, TnpBmax, TnpB_CV2 and TnpBmax_CV2 combined with Trim2 ωRNA scaffold in HEK293T cells at indicated endogenous sites. HDR and NHEJ were measured with ampliconNGS (*n* = 3 biological replicates). Each dot represents an individual biological replicate, and bars represent the mean ± standard deviation. All *P* values were calculated using an unpaired, two-sided *t*-test. ns, not significant (P ≥ 0.05), \**P*L<L0.05, \*\**P*L<L0.01, \*\*\**P*L<L0.001.

**Figure S6.**
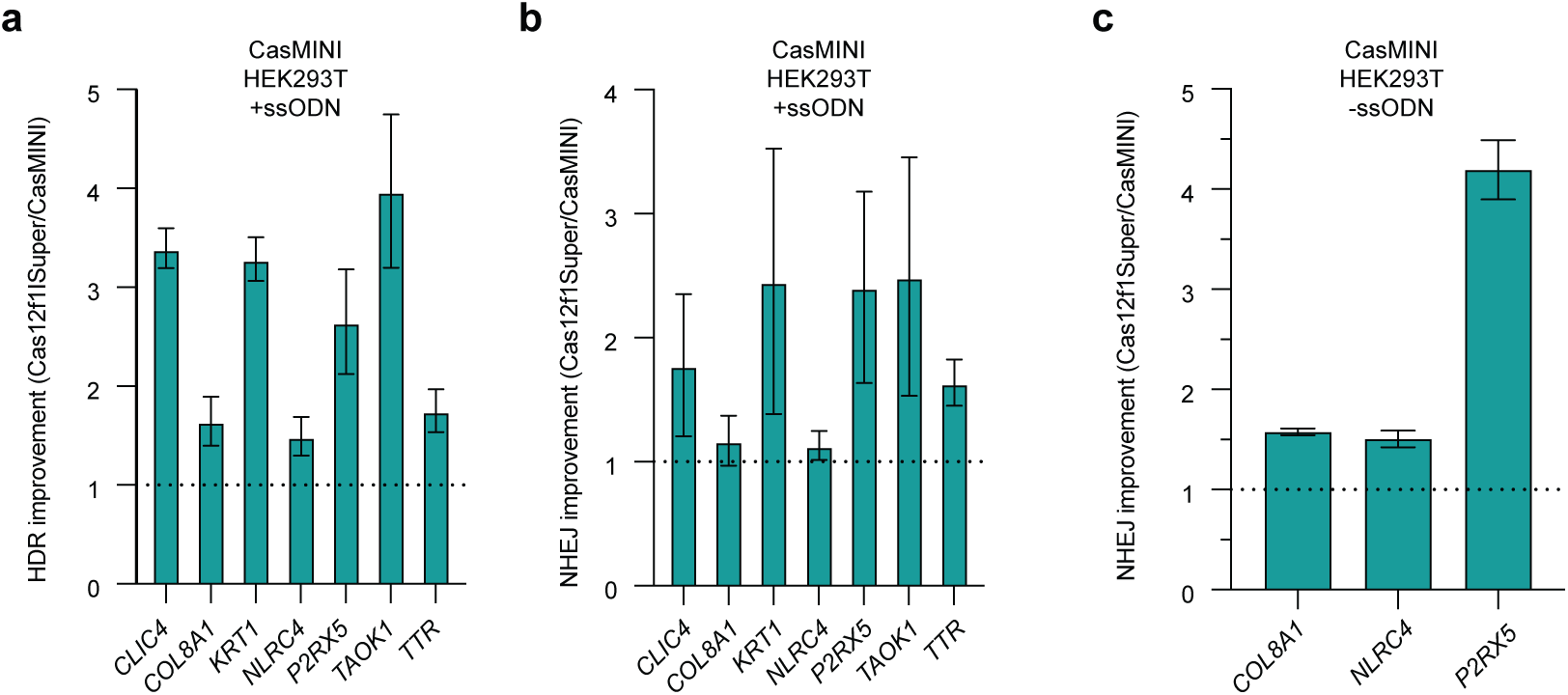
Improved variant Cas12f1Super edits endogenous genomic loci. (**a,b**) Fold change of HDR efficiency (**a**) or NHEJ efficiency (**b**) of Cas12f1Super relative to CasMINI in presence of ssODN measured with ampliconNGS at indicated endogenous loci in HEK293T cells (*n* = 3 biological replicates). (**c**) Fold change of NHEJ efficiency of Cas12f1Super relative to CasMINI in absence of ssODN measured with ampliconNGS at indicated endogenous loci in HEK293T cells (*n* = 3 biological replicates). Bars represent mean ± standard deviation. Fold change was calculated from group means, with propagated standard deviation shown as error bars.

**Figure S7.**
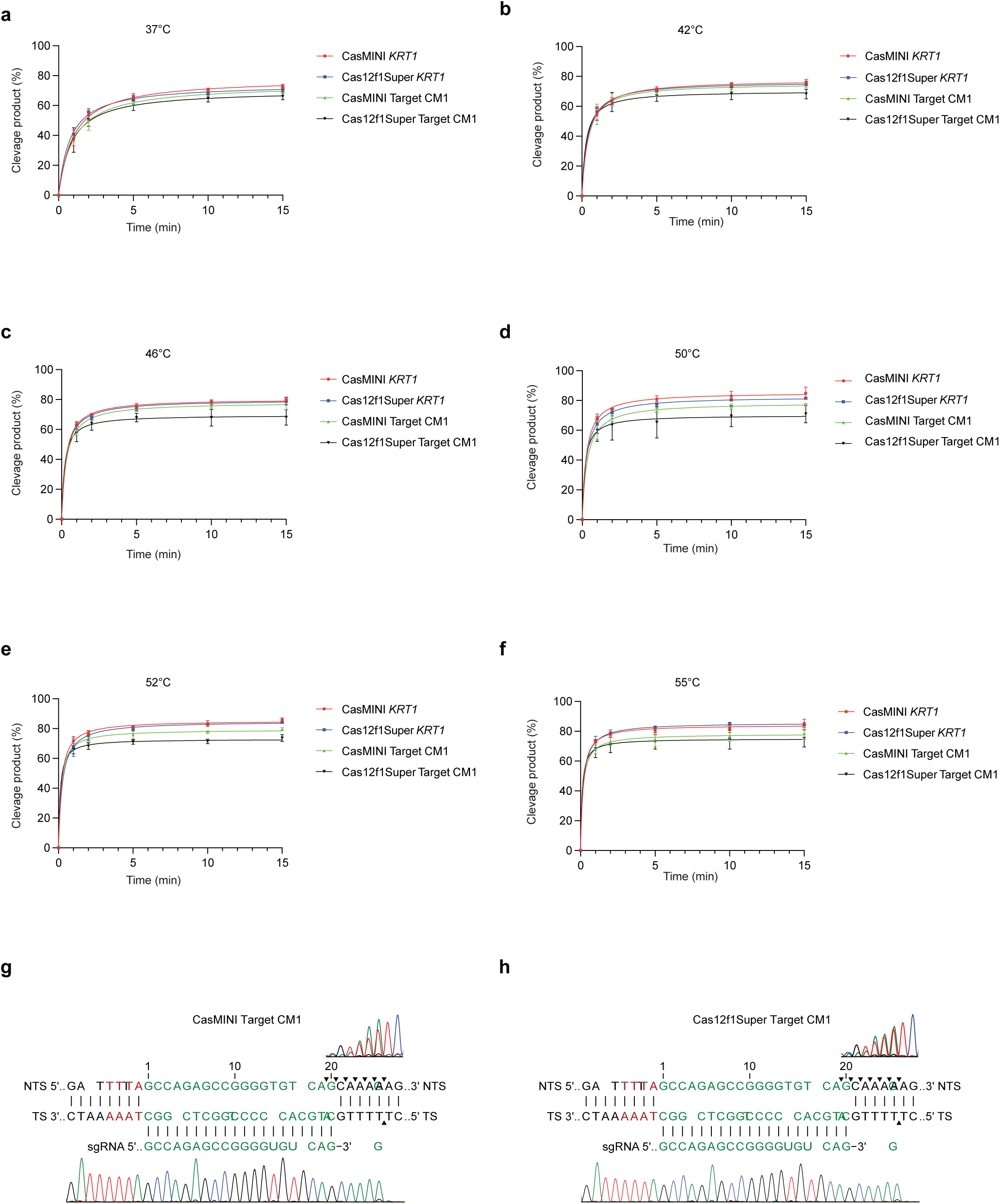
*In vitro* cleavage assays with CasMINI and Cas12f1Super. (**a-f**) Percentages of fully cleaved substrates at two different targets at designated time points by CasMINI or Cas12f1Super (*n* = 3 biological replicates). Each dot represents the mean of three biological replicates ± standard deviation. (**g,h**) Run-off Sanger sequencing of the plasmid products cleaved by CasMINI (**g**) or Cas12f1Super (**h**). Cleavage positions at both the non-targeted strand (NTS) and the targeted strand (TS) are marked with black triangles.

**Figure S8.**
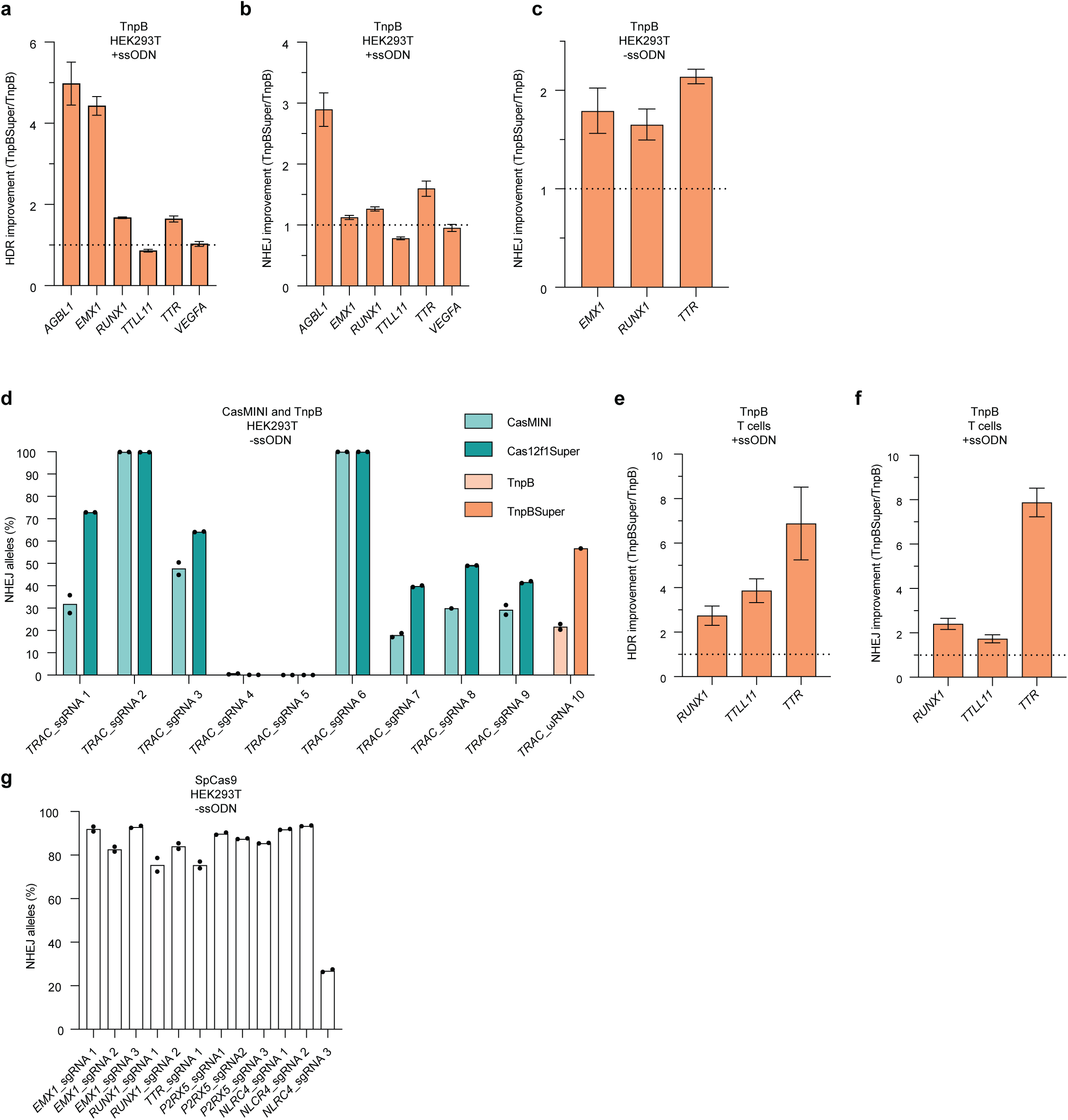
Improved variants Cas12f1Super and TnpBSuper edit endogenous genomic loci. (**a,b**) Fold change of HDR efficiency (**a**) or NHEJ efficiency (**b**) of TnpBSuper relative to TnpB in presence of ssODN measured with ampliconNGS at indicated endogenous loci in HEK293T cells (*n* = 3 biological replicates). (**c**) Fold change of NHEJ efficiency of TnpBSuper relative to TnpB in absence of ssODN measured with ampliconNGS at indicated endogenous loci in HEK293T cells (*n* = 3 biological replicates). Bars represent mean ± standard deviation. Fold change was calculated from group means, with propagated standard deviation shown as error bars. (**d**) Absolute NHEJ efficiency of Cas12f1Super or TnpBSuper measured with ampliconNGS at indicated guides targeting *TRAC* locus in HEK293T cells compared to CasMINI or TnpB (*n* = 2 biological replicates). Each dot represents an individual biological replicate, and bars represent the mean. (**e,f**) Fold change of HDR (**c**) or NHEJ efficiency (**d**) of TnpBSuper relative to TnpB in the presence of ssODN measured with ampliconNGS at indicated endogenous loci in primary T cells (*n* = 3 biological replicates). TnpBSuper or TnpB were electroporated as mRNA, along with synthetic ωRNA and ssODN. (**g**) Absolute NHEJ efficiency of SpCas9 in the absence of ssODN measured with ampliconNGS at indicated at indicated loci in HEK293T cells (*n* = 2 biological replicates). Each dot represents an individual biological replicate, and bars represent the mean.

**Figure S9.**
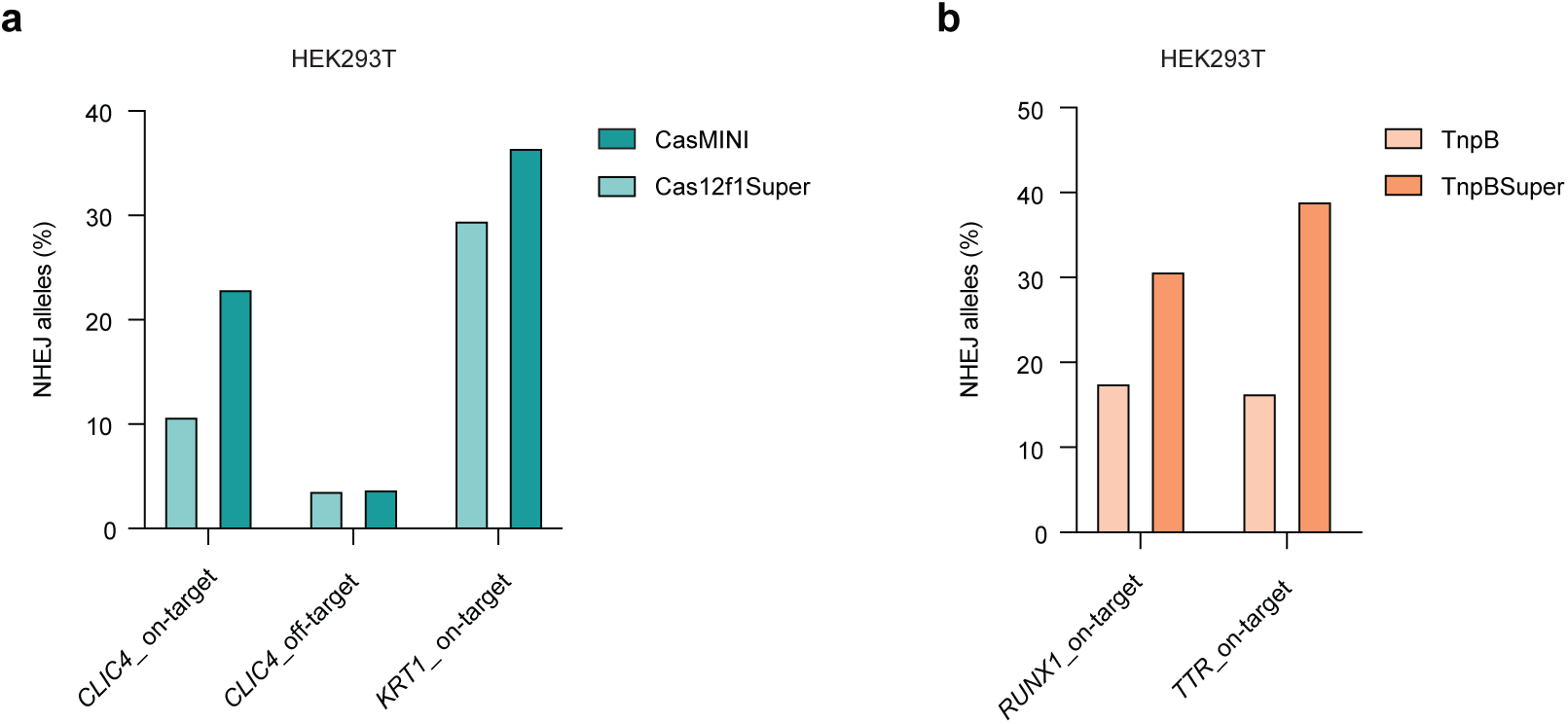
CasMINI/Cas12f1Super and TnpB/TnpBSuper NHEJ efficiency at on-target and off-target sites. (**a**) NHEJ efficiency of CasMINI and Cas12f1Super measured with ampliconNGS at the indicated on-target and off-target sites in HEK293T cells. (**b**) NHEJ efficiency of TnpB and TnpBSuper measured with ampliconNGS at the indicated on-target sites in HEK293T cells. Bars represent the value of a single experimental replicate.

**Figure S10.**
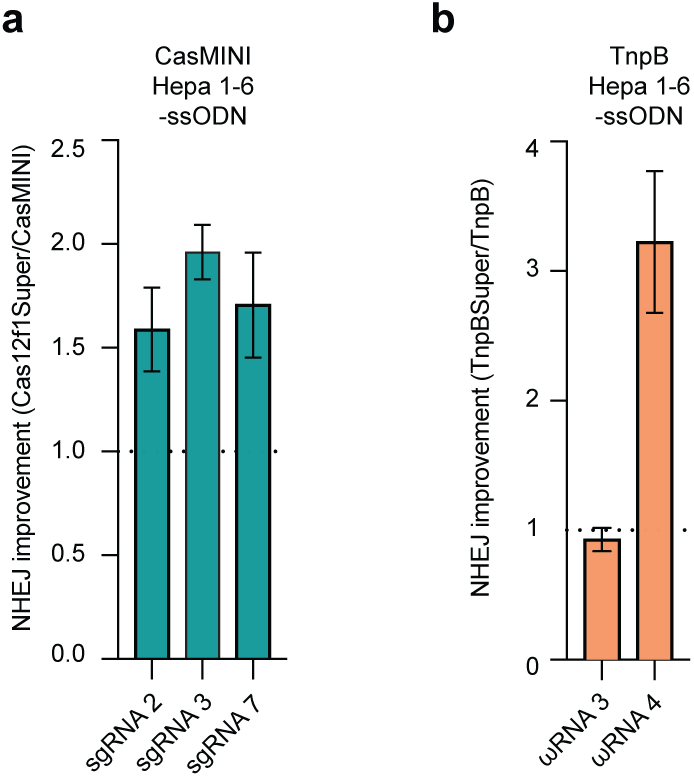
Improved variants Cas12f1Super and TnpBSuper edit endogenous genomic loci. (**a,b**) Fold change of NHEJ efficiency of Cas12f1Super relative to CasMINI (**a**) or of TnpBSuper relative to TnpB (**b**) measured with ampliconNGS at indicated targets within the *Pcsk9* gene in Hepa 1-6 cells (*n* = 3 biological replicates). Each nuclease-guide RNA combination was delivered on a “all-in-one” plasmid. Bars represent mean ± standard deviation. Fold change was calculated from group means, with propagated standard deviation shown as error bars.

**Figure S11.**
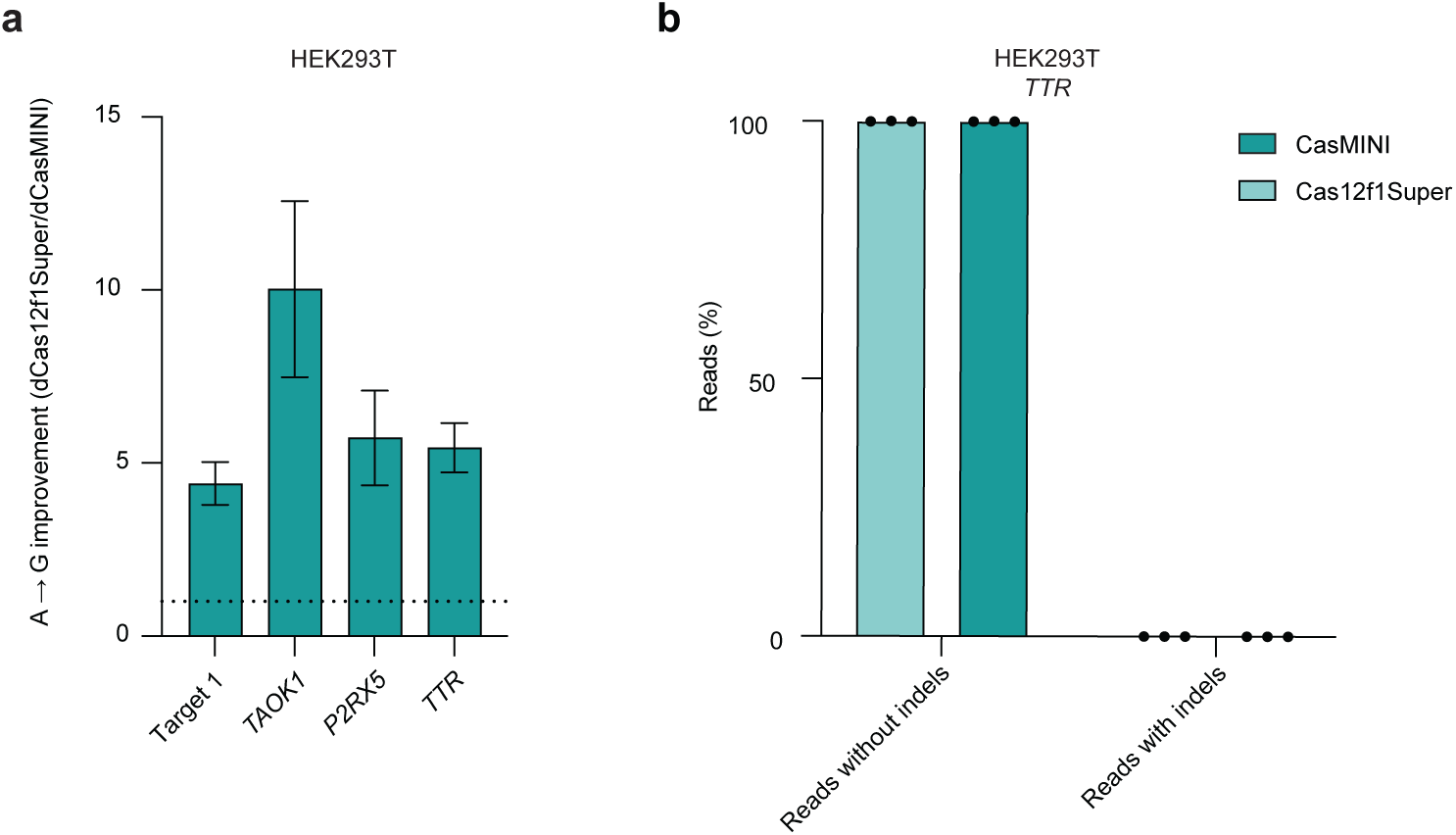
Improved variant dCas12f1Super outperforms dCasMINI as an adenine base editor. (**a**) Fold change of overall adenine base editing activity of dCas12f1Super relative to dCasMINI measured with ampliconNGS at Target CM1 and indicated endogenous sites in HEK293T cells (*n* = 3 biological replicates). Bars represent mean ± standard deviation. Fold change was calculated from group means, with propagated standard deviation shown as error bars. (**b**) Frequency or reads without and with indels upon editing *TTR* site with adenine base editor based on dCasMINI or dCas12f1Super in HEK293T cells (*n* = 3 biological replicates). Each dot represents an individual biological replicate, and bars represent the mean.

**Figure S12.**
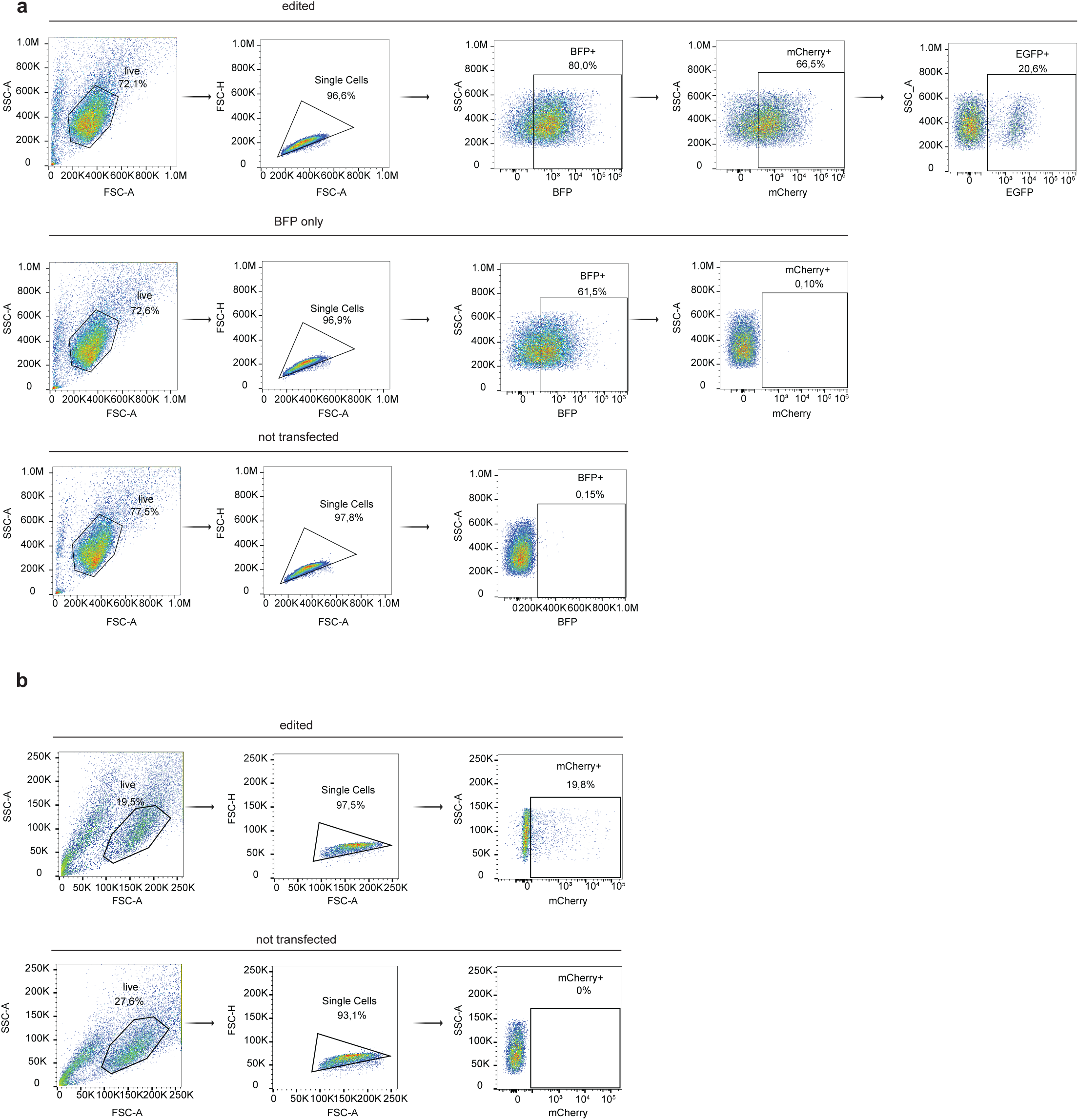
Gating strategies for flow cytometry and FACS experiments related to testing editing at HDR reporter targets and endogenous sites in HEK293T and Hepa 1-6 cell lines. **(a)** Representative plots showing the gating strategy for measuring HDR at HDR reporter targets in HEK293T cell line. Cells were gated for live cells, singlets, BFP positivity, mCherry positivity (BFP marks the guide-expressing construct; mCherry marks the nuclease-expressing construct) and finally, for EGFP positivity. In case of “edited” sample one of the CasMINI variants was co-delivered with sgRNA for Target CM1 together with ssODN. “BFP” only and “not transfected” samples serve as controls for gating. In case of “BFP only” sample only sgRNA targeting Target CM1 was delivered. **(b)** Representative plots showing the gating strategy for FACS-sorting HEK293T or Hepa 1-6 cells for measuring editing. Cells were gated for live cells, singlets and mCherry positivity. In case of “edited” sample CasMINI and sgRNA were delivered on the same “all-in-one” plasmid. “Not transfected” sample serves as control for gating.

**Figure S13.**
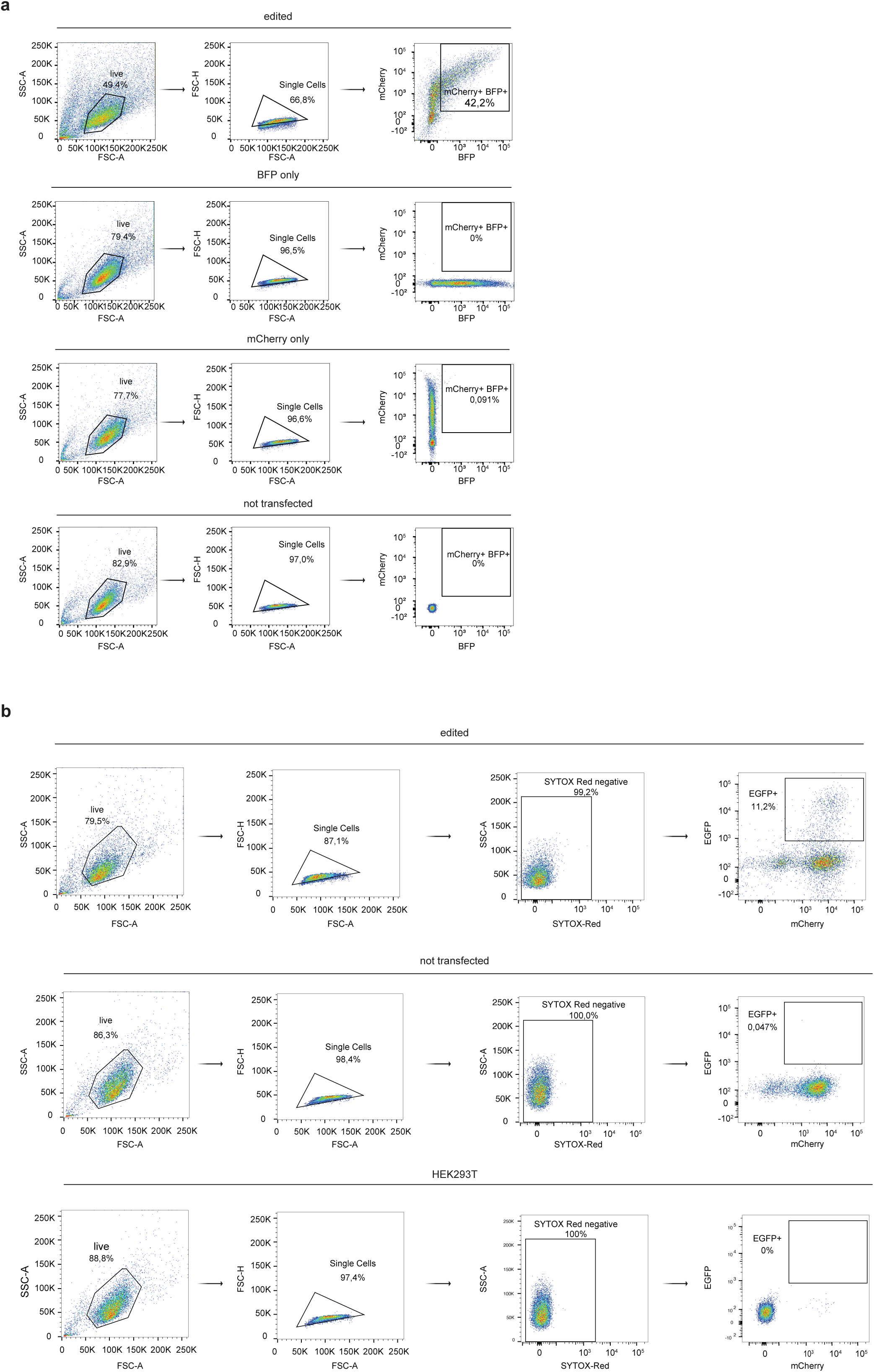
Gating strategy for FACS experiments related to testing editing at endogenous sites in HEK293T and NIH-3T3 cell line and to big scale selection of CasMINI and TnpB for HDR in HEK293T cell line. **(a)** Representative plots showing the gating strategy for FACS-sorting HEK293T and NIH-3T3 cell lines for measuring NHEJ at endogenous sites. Cells were gated for live cells, singlets and mCherry/BFP positivity (BFP marks the guide-expressing construct; mCherry marks the nuclease-expressing construct). In case of “edited” sample CasMINI was co-delivered with sgRNA and ssODN in HEK293T cells. “BFP only”, “mCherry only” and “not transfected” samples serve as controls for gating. **(b)** Representative plots showing the gating strategy for FACS-sorting during round 1 of HDR selection of CasMINI in HEK293T-reporter cell line. Cells were gated for live cells, singlets, SYTOX^TM^-Red-negativity, and finally for mCherry/EGFP positivity. In case of “edited” sample sgRNA for Target CM1 was delivered together with ssODN. “Not transfected” and “HEK293T” samples serve as controls for gating. In case of “not transfected” control sgRNA and ssODN were not delivered.

## Supplementary Tables

**Supplementary Table 1.** Table listing all CasMINI and TnpB evolved variants and combinatorial variants including their mutations.

**Supplementary Table 2.** Table listing the sequences of CasMINI and TnpB reporter and endogenous targets, and all primers and oligonucleotides used in this study.

**Supplementary Table 3.** Table listing all CasMINI and TnpB ssODNs used for reporter and endogenous targets.

**Supplementary Table 4.** Table listing all plasmids used in this study.

**Supplementary Table 5.** Table listing sgRNA sequences used *in vitro* cleavage assays for CasMINI and Cas12f1Super.

**Supplementary Table 6.** Table listing all CasMINI and TnpB guide RNA scaffolds used in this study.

## Notes

### Summary of Updates

We have revised the manuscript and performed additional experiments to address the following: editing at endogenous sites in the absence of a homology donor, benchmarking of Cas12f1Super and TnpBSuper systems versus SpCas9, and evaluation in the therapeutic context of in vivo editing using AAV. We also clarified that our selection strategy enriches for overall editing activity rather than HDR alone.

